# Comprehensive Atomic-Scale 3D Viral-Host Protein Interactomes Enable Dissection of Key Mechanisms and Evolutionary Processes Underlying Viral Pathogenesis

**DOI:** 10.1101/2025.03.28.645946

**Authors:** Le Li, Priyamvada Guha Roy, Yilin Liu, Zizhao Zhang, Dapeng Xiong, Ram Savan, Nandan S. Gokhale, Luis M Schang, Jishnu Das, Haiyuan Yu

## Abstract

Viral-human protein interactions are critical for viral replication and modulation of the host immune response. Structural modeling of these interactions is vital for developing effective antiviral therapies and vaccines. However, 99% of experimentally determined binary host-viral interactions currently lack structural information. We aimed to address this gap by leveraging computational protein structure prediction methods. Using extensive benchmarking, we found AlphaFold to be the most accurate structure prediction model for host-pathogen protein interactions. We then predicted the structures of 11,666 binary protein interactions across 33 viral families and created the most comprehensive atomic-scale 3D viral-host protein interactomes till date (https://3d-viralhuman.yulab.org).

By integrating these interactomes with genetic variation data, we identified population-specific signatures of selection on variants coding for interfaces of viral-human interactions. We also found that viral interaction interfaces were less conserved than non-interface regions, a striking trend that is opposite to what is observed for host interfaces, suggesting different evolutionary pressures. Systematic analyses of interface sharing between host and viral proteins binding to the same host protein revealed mutation rate-dependent differences in interface mimicry. Similar mutation rate-dependent differences were seen in the interface sharing between viral proteins binding to a host protein. We also found that the patterns of E6 protein binding to KPNA2 differed between high– and low-risk oncogenic human papillomaviruses (HPVs), and clustering based on these binding patterns allowed the classification of HPVs with unknown oncogenic risk. Our interface mimicry analyses also unveiled a novel mechanism by which herpes simplex virus-1 UL37 suppresses the antiviral immune response through disruption of the TRAF6-MAVS signalosome interaction.

Overall, our comprehensive 3D viral interactomes provide a resource at unprecedented scale and resolution that will enable researchers to explore how variation and signatures of selection influence viral interactions and disease progression. This tool also facilitates the identification of conserved and unique interaction patterns across viruses, empowering researchers to generate testable hypotheses and ultimately accelerate the discovery of novel therapeutic targets and intervention strategies.

## Introduction

As obligate parasites, viruses depend on human cellular proteins—including for entry, transcription, translation, metabolism, genome replication, trafficking, and egress—for their replication, relying on a complex network of protein-protein interactions to accomplish this goal (Lasso et al. 2019). Interactions between viral and host proteins also play a key role in modulating innate immune responses. Studying the structure and dynamics of these interactions can reveal new targets for antiviral therapies and vaccines and help develop strategies to bolster host defenses (de Chassey et al. 2014; Cakir et al. 2021; Wierbowski et al. 2021). For instance, the discovery of CCR5 as a crucial co-receptor for HIV1 attachment led to the development of maraviroc, a CCR5 antagonist with strong anti-HIV activity (Fätkenheuer et al. 2005). Moreover, elucidating the binding interface between the SARS-CoV-2 spike protein and the human ACE2 receptor has been instrumental in the development of therapeutic agents targeting this interaction (Lan et al. 2020; Ferrari et al. 2021; Shoemaker et al. 2022). Integrating structural information with genetic data can uncover key variants that modulate the immune cascade and systemic response to viral infections (Chhibbar et al. 2024; Wierbowski et al. 2021). Despite their importance, however, structural data for most viral-host protein interactions remain scarce due to the challenges of experimentally resolving complex structures. This gap underscores the need for comprehensive 3D modeling of viral-human protein interactions, which could provide key molecular insights into viral pathogenesis and inform the development of targeted antiviral strategies.

Computational methods have been widely used to infer structural information for protein interactions, from predicting interaction interfaces to modeling entire protein complexes (Xiong et al. 2024; Abramson et al. 2024; Bryant and Noé 2023; Lin et al. 2023; Ketata et al. 2023; Dominguez, Boelens, and Bonvin 2003)(Jumper et al. 2021; Abramson et al. 2024; Evans et al. 2022);(Xiong et al. 2024; Abramson et al. 2024; Bryant and Noé 2023; Lin et al. 2023; Ketata et al. 2023; Dominguez, Boelens, and Bonvin 2003). We evaluated the ability of state-of-the-art methods, including AlphaFold (Jumper et al. 2021; Abramson et al. 2024; Evans et al. 2022), ESMFold (Lin et al. 2023), DiffDock-PP (Ketata et al. 2023), HADDOCK (Dominguez, Boelens, and Bonvin 2003), and ECLAIR+HADDOCK (Wierbowski et al. 2021), in generating structural models for host-viral protein interactions using a comprehensive benchmark dataset of 509 host-pathogen protein interactions with PDB structures (Wierbowski et al. 2021). Despite relying on paired multiple sequence alignments (MSAs), which are largely incomplete for host-viral protein interactions due to the rapid evolution of viral proteins, lack of homologous sequences, and asymmetric co-evolution between host and viral proteins, AlphaFold had the best performance in terms of predicting overall structures and specific interface residues. Therefore, we used AlphaFold to predict the co-complex structures of all biochemically verified viral-human protein interactions in the literature to date, creating the most comprehensive 3D viral-human interactomes to date, spanning 11,666 direct, binary protein interactions across 33 viral families.

We leveraged this dataset to examine the evolutionary dynamics of host-virus interactions and identified population-specific selective pressures acting on host interface residues. Further, our analysis revealed that viral interface residues exhibited a distinct pattern of conservation compared to non-interface residues within viral proteins. Moreover, the difference in evolutionary constraints between viral interface and non-interface residues followed a pattern fundamentally different from what was observed in endogenous host-host protein interactions, indicating differential selective pressures. We examined the extent of interface sharing between host-host and host-virus protein interactions and found that, across viral families, mutation rate largely dictated patterns of interface mimicry. As a validation, the binding interfaces of E6 protein from human papillomaviruses (HPVs) on different human proteins served as a distinguishing factor between non-oncogenic and high– and low-risk oncogenic types. As another independent validation, we identified a novel mechanism through which herpes simplex virus-1 (HSV1) UL37 inhibits the antiviral immune response by interrupting the interaction between TRAF6 and MAVS.

Our compendium of 3D viral interactomes is a powerful resource for studying the dynamics and structural basis of host-viral interactions. It provides a unique platform for integrating multi-omic data to examine how genetic and proteomic variations in pro– and anti-viral host factors influence interactions with viral components and modulate downstream immune activation. This comprehensive framework not only enables the identification of key interaction patterns but also empowers researchers to generate testable mechanistic hypotheses, driving the discovery of novel therapeutic targets and intervention strategies.

## Results

### 3D Viral-Human Protein Complex Structural Modeling

Understanding the structural basis of viral-human protein interactions is critical for uncovering viral replication mechanisms and may help to identify novel therapeutic targets. Due to experimental challenges, however, structural data for these interactions remain scarce. Computational modeling provides a promising alternative for generating 3D structural models of viral-human complexes, enabling large-scale investigations of protein interactions at an atomic resolution.

To identify the most reliable computational approach for modeling viral-human protein interactions, we systematically benchmarked state-of-the-art protein structure prediction methods, including AlphaFold, ESMFold (Lin et al. 2023), DiffDock-PP (Ketata et al. 2023), HADDOCK (Dominguez, Boelens, and Bonvin 2003), and ECLAIR+HADDOCK (Wierbowski et al. 2021).

We first evaluated the accuracy of overall structure prediction of these methods using our benchmark set of 509 pathogen-host protein interactions with PDB structures (Wierbowski et al. 2021). AlphaFold outperformed all other methods in terms of DockQ score, interface root mean square deviation (iRMSD), and ligand RMSD (lRMSD) (Figures 1A-1C). Specifically, over 25% of AlphaFold predictions were classified as high quality (DockQ score ≥ 0.8, green portion in Figure 1A), while approximately 50% fell within the medium to high range (DockQ ≥ 0.49, yellow portion in Figure 1A). In contrast, the best alternative method produced fewer than 5% high quality predictions and less than 10% acceptable predictions (DockQ ≥ 0.23). Additionally, AlphaFold achieved significantly lower iRMSD (<5 Å vs. ∼15 Å) and lRMSD (<10 Å vs. >30 Å) compared to other methods, underscoring its superior structural accuracy.

**Figure 1.**
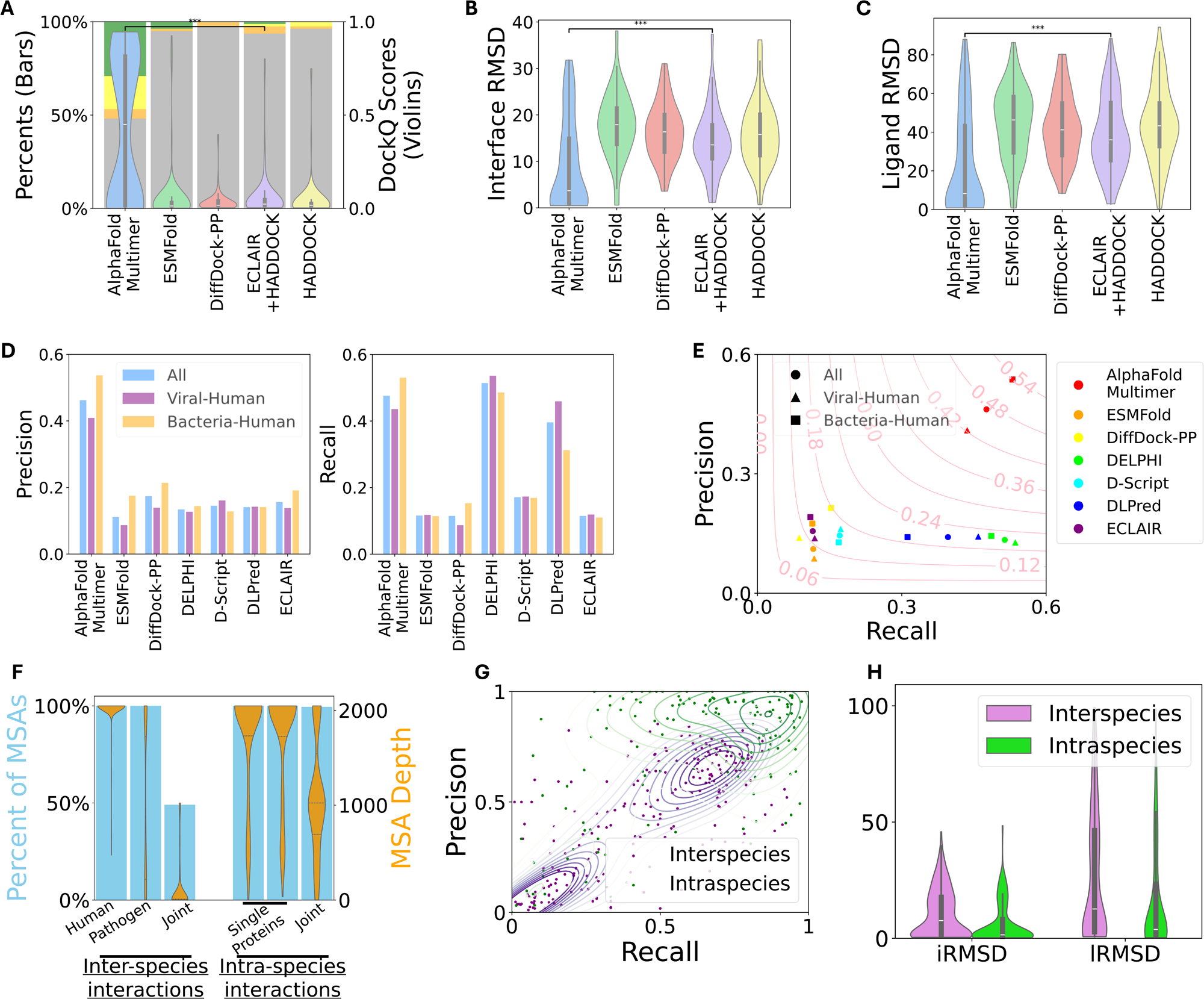
Benchmark of 3D Structure Prediction of Pathogen-Host Protein Interactions. (A)-(C) Performance of computational approaches in predicting structures for pathogen-host interactions, evaluated by DockQ (A), iRMSD (B), and lRMSD (C). Prediction quality based on DockQ scores is categorized as: Incorrect (DockQ < 0.23), Acceptable (0.23 ≤ DockQ < 0.49), Medium (0.49 ≤ DockQ < 0.8), and High (DockQ ≥ 0.8). Significance levels of the differences between AlphaFold and the best alternative methods are indicated as follows: ‘ns’ – not significant, ‘*’ – p-value < 0.05, ‘**’ – p-value < 0.01, ‘***’ – p-value < 0.001. (D)-(E) Performance of computational approaches in predicting interface residues for pathogen-host protein interactions in terms of macro precision and recall (a single precision and recall are calculated from the whole set of predicted interface residues for all interactions). The performance is shown for all interactions as well as for viral-human and bacteria-human subsets. (F) Depth of MSAs for inter-species and intra-species interactions. The bars display the percentage of interactions with individual and joint MSAs, while the violin plots show the distributions of MSA depths. (G) Scatter plots showing precision and recall of predicted interface residues for individual interactions. Contour lines indicate the density of predictions for both inter-species and intra-species interactions. (H) Comparison of structure-level quality measurements, including iRMSD and ligand RMSD lRMSD, between inter-species and intra-species interactions.

Accurately predicting binding interfaces is key to modeling protein interactions. Therefore, we specifically evaluated these tools for their performance in predicting experimentally known interface residues. We further included state-of-the-art interface prediction tools including DELPHI, D-Script, DLPred, and ECLAIR, together with the structure prediction methods mentioned above. Our pathogen-host benchmark set contains both bacterial and viral pathogens. We further divided the benchmark set into bacterial-host and viral-host subsets.

Across the whole benchmark set and viral-human and bacterial-human subsets, AlphaFold exhibited the highest precision (>0.4 vs. <0.2 for the best alternative method, left panel of Figure 1D) while maintaining a recall close to the best-performing method (∼0.5, right panel Figure 1D). The resulting F1 score of AlphaFold (0.42) significantly outperformed other tools (≤0.24, Figure 1E), further reinforcing its superior reliability in interface prediction. Notably, bacterial-human interactions were modeled more accurately than viral-human interactions, which aligns with observation that higher mutation rates and genetic diversity among viruses than bacterial result in lower-quality MSA (Alqahtani and Almutairy 2023) (Supplementary Figure 1A).

Given that MSAs play a crucial role in AlphaFold-based predictions, we first examined the impact of MSA depth on inter-species (pathogen-host) interactions compared to intra-species interactions. We found that while most intra-species interactions had a high-depth of joint MSAs (median depth ∼1000), over 50% of pathogen-host interactions lacked joint MSAs, and most of the remaining ones had low coverage (<100 sequences per protein) (Figure 1F). This disparity directly affected prediction quality, as interface precision, recall, and structural similarity metrics (iRMSD and lRMSD) were consistently higher for intra-species interactions than for pathogen-host interactions (Figures 1G-1H), which has been reported previously (Zhu et al. 2023; Lupo, Sgarbossa, and Bitbol 2024; Tsuchiya, Yamamori, and Tomii 2022). Similar results were obtained for the subsets of novel interactions (not used in training of AlphaFold) only (Supplementary Figure 1B-C)

Given the recent development of AlphaFold3 (Abramson et al. 2024), we also directly compared its performance with AlphaFold-Multimer (AFM) on viral-human interactions with available PDB structures (May 2024). To ensure a rigorous evaluation, we focused only on heterodimers corresponding to which there were known interfaces based on co-crystal structures. The results (Supplementary Figure 1D) indicate that AlphaFold3 exhibited higher recall, but lower precision for interface residue prediction compared to AFM. Therefore, we decided to apply AFM (henceforth referred to as AlphaFold) for large-scale viral-human interaction modeling.

In summary, despite the challenges of low-coverage joint MSAs for inter-species interactions, we find that AlphaFold is currently the best tool for accurate structure prediction, including precise interface predictions, significantly outperforming other state-of-the-art methods. Therefore, we applied AlphaFold to construct comprehensive 3D viral-human interactomes (Methods).

### Creating Comprehensive 3D Viral-Human Interactomes

We collected viral-human protein-protein interaction data from VirHostNet 2.0, IntAct (release 2022.02.03), and BioGRID (version 4.4.209). After mapping proteins to UniProt IDs and retrieving taxonomy information, we obtained 65,823 interactions. Based on experimental evidence, interactions were categorized as binary (direct interactions, 14,845) or co-complex (membership in the same complex, 51,866). This study focuses exclusively on the 11,666 binary viral-human interactions remaining after filtering. Of the 11,666 viral-human and 2,838 viral-viral biochemically validated binary protein interactions we curated, approximately 99% and 95% of the viral-human and viral-viral interactions, respectively, did not have a corresponding structure (Figure 2A). This highlights the major gap in the availability of structural data for viral protein interactions which is critical for understanding the molecular basis of disease pathogenesis.

**Figure 2.**
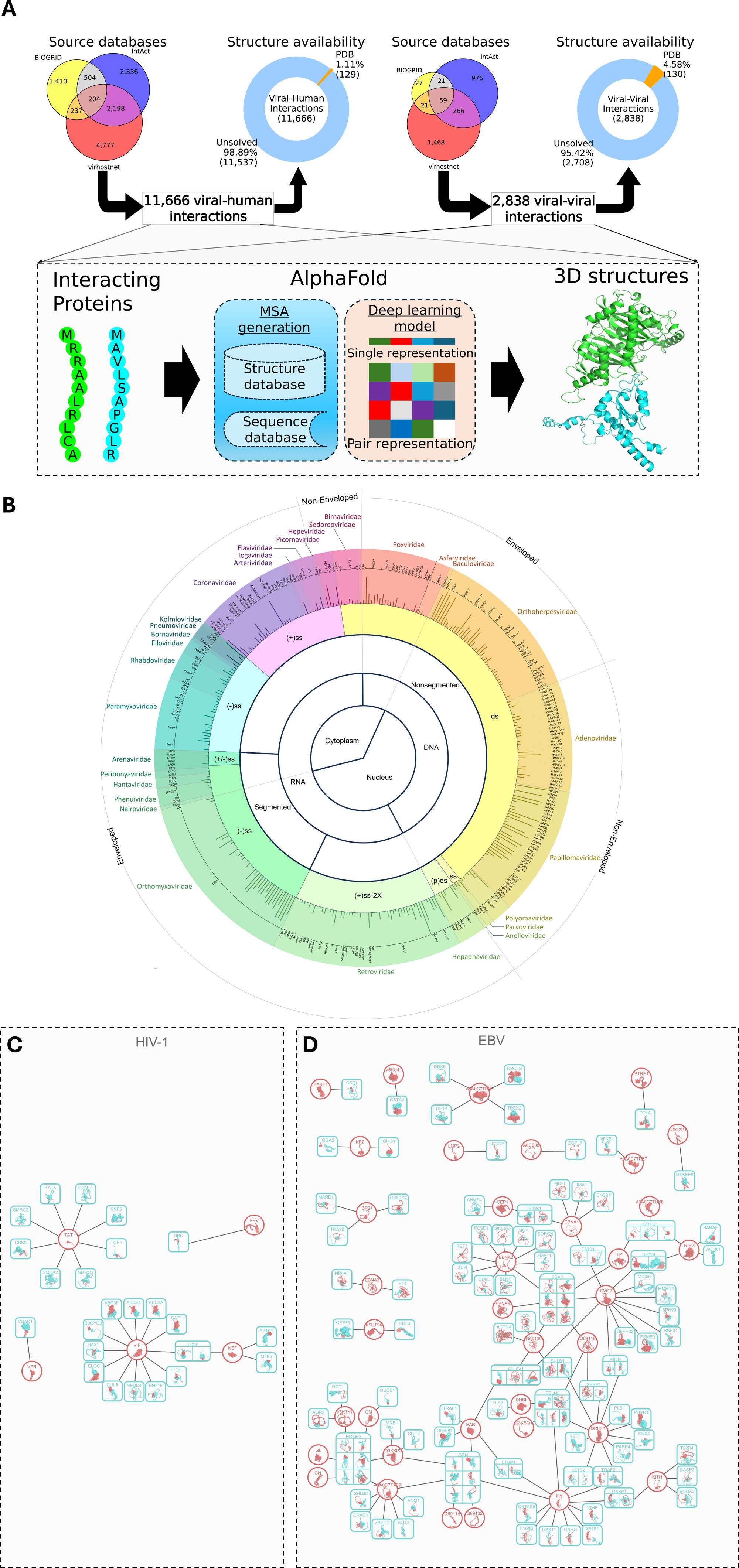
Creating Comprehensive 3D Viral-Human Interactomes. (A) We collected a total of 11,666 binary viral-human interactions and 2,838 viral-viral interactions from three public databases: IntAct, BIOGRID, and VirHostNet, as of May 2022. Among these interactions, 1.11% (129) of the viral-human protein interactions and 4.58% (130) of the viral-viral protein interactions have known 3D structures (full or partial) available in the PDB database. The structures of the known interacting protein pairs without known 3D structures were predicted using AlphaFold. (B) A circular plot illustrating the architecture of all viruses included in this study. The viruses are classified in multiple tracks based on multiple categories (from inner to outer): nucleus vs. cytoplasm (the first track), DNA vs. RNA (the second track), segmented vs. nonsegmented (the third track), and single-strand vs. double-strand (the fourth track, ss: single-strand, ds: double-strand, (p)ds: partial double-strand, (-): negative sense, (+): positive sense, (+/-): ambisense, 2X: two copies). The outermost track indicates the common name and the number of interactions (represented by bars) for each virus, grouped and colored according to their family names (outer labels), which are further labeled based on whether they are enveloped or non-enveloped. (C) The known viral-human protein interactome for HIV-1 group M subtype B (isolate PCV12). (D) The known viral-human protein interactome for EBV strain GD1. In both (C) and (D), viral proteins are shown in red and human proteins in cyan. Circles represent single viral proteins, while rectangles represent viral-human protein complexes.

To this end, we predicted the co-complex structures of all collected binary viral-human and viral-viral interactions using AlphaFold (Methods; Figure 2A). The resulting dataset includes 360 viruses, across 31 families, which are visualized in a circular plot based on cellular site of replication, genetic materials (DNA or RNA, segmented or not, genome sense and single or double stranded), family, and whether they are enveloped or not (Figure 2B).

This comprehensive collection of 3D viral interactomes serves as a powerful resource for generating and testing hypotheses for single-virus studies or large-scale viral studies. Unlike a traditional viral-human interactome that lacks structural information, a 3D interactome provides structural information that reveal key functional insights (e.g. binding sites and pockets), which allows for a precise understanding of interaction mechanisms, identification of potential therapeutic targets, and development of structure-based drug design strategies. The 3D viral-human interactomes of HIV-1 group M subtype B/isolate PCV12 and Epstein-Barr virus (EBV) strain GD1 are visualized in Figures 2C and 2D, respectively.

### Accurate Predictions of Previously Solved Structures and Biochemically Validated Interface Residues

Among the 11,666 viral-human interactions, 1.11% (129 interactions) had PDB structures, which were included in the benchmark to validate our interface predictions. The average precision and recall of the predicted interfaces were approximately 0.5 and 0.45, respectively, with over half of the interactions exhibiting good precision and recall over 0.5 (Figure 1B). To further test whether AlphaFold was able to learn previously unseen structures, we explored its performance on 25 interactions that were previously unseen by AlphaFold and observed a similar distribution and values of precision and recall (0.48 and 0.42, respectively, Supplementary Figure 2).

When the predicted and known PDB structures for the benchmark set were compared, the predictions closely matched the PDB structures. For the SARS-CoV-2 NS6 and RAEL 1 interaction (NS6_SARS2 and RAE1L_HUMAN, respectively), the known PDB structure (7vph, released on January 19, 2022) covered only a 21 amino acid segment of NS6. AlphaFold perfectly predicted the existing structures (TM-score: 0.99, RMSD: 0.47) and showed reasonable performance for the unsolved viral protein segments (shown in red in Figure 3A), despite the lack of a joint MSA. AlphaFold also accurately modeled the the known PDB structure (8cx0, released on February 15, 2023) for the interaction between HIV-1 Vif and PEBB (VIF_HV1H2 and PEBB_HUMAN, respectively, TM-score: 0.83, RMSD: 1.87), despite a low-depth MSA (99 sequences) for Vif and a joint MSA depth of 2, and predicted the structures of the unsolved segment of PEBB (shown in pink in Figure 3B). Similar performance was observed for the predictions of Yaba-like disease virus 16L and B2L11 interaction (16L_YLDV and B2L11_HUMAN, respectively, 6tqq, TM-score: 0.83, RMSD: 1.17 Å, Figure 3C) and Cedar virus protein G and EFNB2 interaction (G_9MONO and EFNB2_HUMAN, respectively, 6p7y, TM-score: 0.83, RMSD: 1.7 Å, Figure 3D). The high concordance between the predictions and known structures suggested that the performance of AlphaFold was generalizable to all other viral-human interactions in our database.

**Figure 3.**
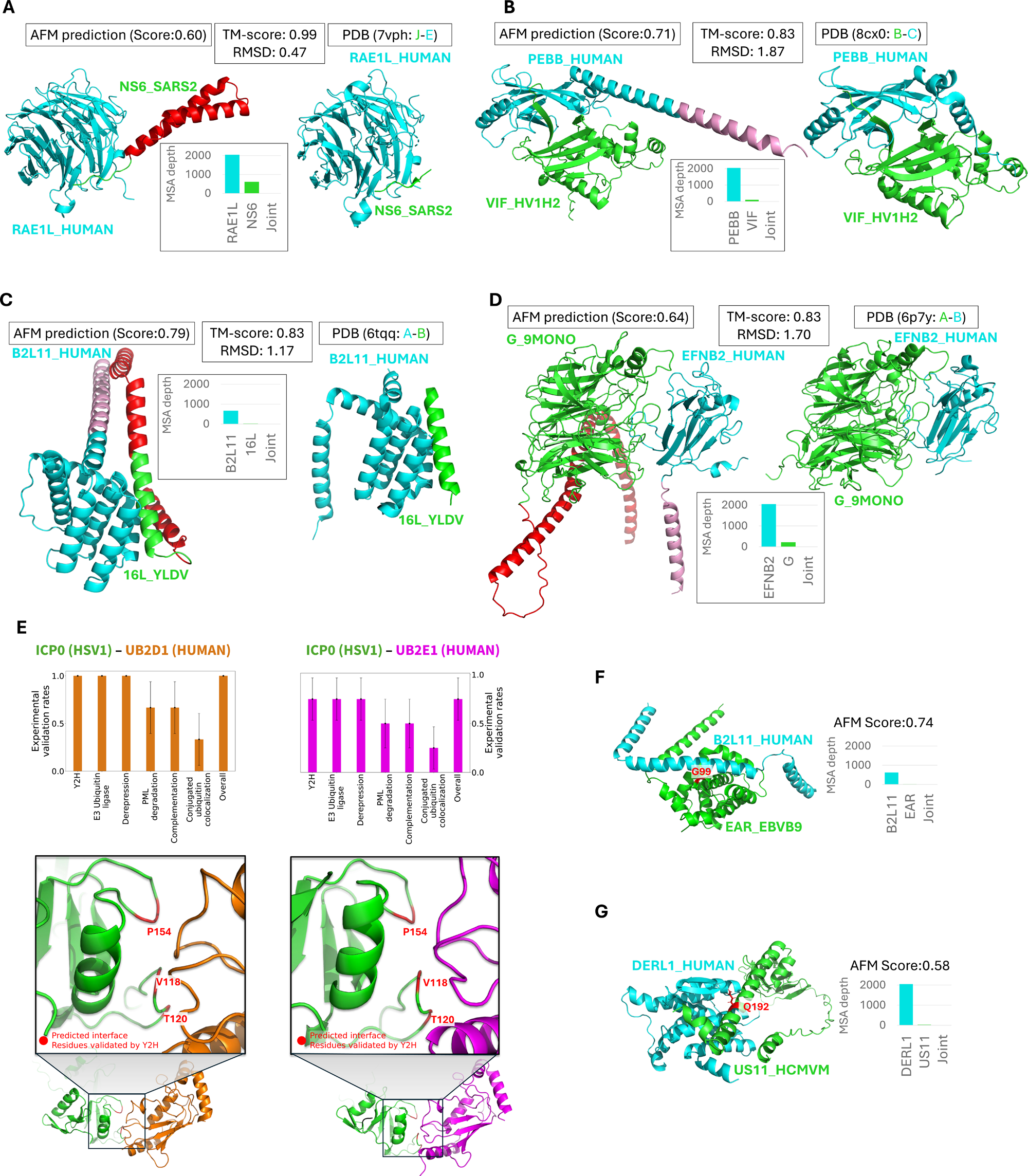
Validation of the 3D Structures of Viral-Human Protein Interactomes Predicted by AlphaFold. (A)-(B) Predicted structures of the interactions between RAE1L and NS6 (SARS2) (A) and PEBB and VIF (HV1H2) (B). The depths of MSAs for individual proteins and joint MSAs are included. Novel structure fragments predicted by AlphaFold are colored pink for human protein regions and red for viral protein regions. (C)-(D) Structures predicted by AlphaFold for the interactions of B2L11 with 16L (YLDV) (C) and EFNB2 with G (9MONO) (D). The depths of MSAs for individual proteins and joint MSAs are included. Novel structure fragments predicted by AlphaFold are colored pink for human protein regions and red for viral protein regions. (E) Predicted structure of the interaction between UB2D1/UB2E1 and ICP0 (HSV1), which has been experimentally validated. Bar plots at the top show the rates of predicted interface residues that were experimentally validated to affect different phenotypes. Zoomed in panels at the bottom indicates the predicted interface residues validated by Y2H and other experiments. (F) Predicted structure of the interaction between B2L11 and EAR (EBVB9) with the experimentally validated interface residue (G99) emphasized. (G) Predicted structure of the interaction between DERL1 and US11 (HCMVM) with experimentally validated interface residue (Q192) emphasized.

Overall, we identified several predicted interaction interfaces that play key roles in mediating the interactions between the proteins that have been validated through mutagenesis studies. One such example includes the structural predictions for the interactions between herpes simplex virus 1 (HSV-1) (or human herpesvirus 1, HHV1) ICP0 protein and human UB2D1 and UB2E1 proteins, interactions which counteract host antiviral defense mechanisms (Smith, Boutell, and Davido 2011). The immediate-early protein ICP0 promotes viral gene expression and replication by leveraging its E3 ubiquitin ligase activity to induce degradation of several cellular defense proteins. ICP0 engages with UBE2D1 and UBE2E1, which are E2 ubiquitin-conjugating enzymes, to target cellular proteins for degradation by the proteasome, creating a more favorable environment for viral gene expression and replication. These interactions allow ICP0 to counteract host cell intrinsic and innate antiviral defenses by inducing degradation of antiviral proteins such as PML and Sp100 (Boutell and Everett 2013). In a double-blinded cross-validation with a comprehensive study investigating twelve point mutations on HSV1 ICP0 that affect its interactions with UB2D1 and UB2E1 and the consequent phenotypes, including E3 ubiquitin ligase activity, conjugated ubiquitin colocalization, promyelocytic leukemia protein degradation, complementation of plaque formation efficiency, and derepression of gene expression (Vanni et al. 2012), we found that our predicted interfaces highly overlapped with the Y2H-validated interfaces (Figure 3E). In the predicted structure of HSV1 ICP0 and UB2D1 complex (left panel of Figure 3E), we identified V118, T120, and P154 as interface residues (SASA > 15 Å^!^, ΔSASA > 1 Å^!^), which influence various interaction-related phenotypes. Similarly, in the predicted structure of ICP0 and UB2E1 complex (right panel of Figure 3E), we identified V118, T120, R150, and P154 as interface residues, and of these, three positions were experimentally verified. The resulting validation rates for the predicted interfaces ranged from 0.75 to 1 (top panel of Figure 3E).

We also found experimental data showing mutagenesis of G99 at the interaction interface of EBV BHRF1 protein (EAR_EBVB9) with B2L11 (B2L11_HUMAN) disrupted binding of the two proteins. BHRF1, an anti-apoptotic protein, binds to B2L11, a pro-apoptotic protein, to manipulate cell death pathways, promoting cell survival and viral persistence (Desbien, Kappler, and Marrack 2009); Figure 3F). The mutant lost this interaction and cells expressing this mutant were no longer protected from apoptotic signals. AlphaFold also successfully predicted the interface residue (Q192) critical for the interaction between Derlin-1 (DERL1_HUMAN) and the human cytomegalovirus (HCMV) protein US11 (US11_HCMV), which has been experimentally validated (Lilley and Ploegh 2004; Xiaoyang Liu et al. 2019), Figure 3G). HCMV US11 exploits the Derlin-1-dependent endoplasmic reticulum-associated protein degradation pathway to degrade major histocompatibility complex class I molecules, allowing the virus to evade immune surveillance (Cho, Lee, and Jun 2013; Cho et al. 2013). We also predicted de-novo the TRAF6 binding motif (_1099_PVEDDE_1101A4_) on the HSV-1 protein UL37 (Xueqiao Liu et al. 2008). Site specific mutation of E1101 significantly impacts binding of these proteins and prevents downstream NF-kappaB activation. Further, we predicted an interface residue (W125) essential for binding of Sendai virus C protein to Alix (Oda et al. 2021), which is crucial for viral budding.

Most of these interactions were predicted using low-quality viral protein MSAs and in the absence of joint MSAs, indicating that AlphaFold was able to infer certain intrinsic patterns from sequence context alone. This highlights the capability of in-silico methods in predicting co-complex structures of viral-human interactions, even when comprehensive MSAs are not available.

### Interface Residues on Human Proteins Show Signatures of Positive Selection

Infections are one of the most significant drivers of selection pressure on the human genome, with patterns of selection shaped by various aspects of host-pathogen interactions—such as exposure duration, geographical spread, morbidity and mortality, and environmental events (Fumagalli et al. 2011; Karlsson, Kwiatkowski, and Sabeti 2014). Differences in these dynamics result in varying selection patterns across populations, resulting in differing burdens of infectious diseases. Host-viral protein interactions are critical for viral replication and propagation. Additionally, certain interactions between viral and host proteins trigger signalling cascades essential for the host immune response against the virus, which are often downmodulated by virus-host protein interactions. The underlying host genome plays a key role in modulating this immune response (Chhibbar et al. 2024). Studying these host-viral protein interactions provides critical insights into differential disease burden for certain infectious diseases.

Given that interaction interfaces play a key role in modulating protein interactions, we hypothesized that these interface regions would be under stronger constraints than other regions of the same proteins. To test this hypothesis, we estimated the fixation indices (F_ST_) for all coding variants within human proteins from our database using the 1000 Genomes data for all population pairs (EUR, AFR, EAS, SAS). Fixation index is a measure of allelic divergence between populations and a large difference in allelic frequency between populations can be an indicator of selection (Wright 1965; Lewontin and Krakauer 1973). These coding variants were then categorized into interface and non-interface variants based on structurally-resolved interactome. Thereafter, we tested the difference in proportion of interface vs non-interface variants with high F_ST_ to identify virus specific signatures of selection in different populations (Figure 4A). Stratifying variants based on whether they were at the interface or not, revealed population-specific patterns of selection for different viruses (Figure 4B). SNPs with high rates of allelic divergence for the South-Asian population appear to colocalize at the interface of protein-protein interactions for a large proportion of the viruses analyzed. Moreover, protein interfaces engaged in interactions with EBV, HPV11, and HPV18 viral proteins showed evidence of selection in multiple populations. When the same analysis was repeated using variants coding for amino-acid residues at any surface of proteins involved in interactions with different viruses, not just those engaged in interactions with the viral proteins, selection was observed only for HPV8 (Figures 4C and 4D). Next, we focused on all residues that are at the surface of these proteins. Since interface residues are a subset of surface residues, this analysis helps test the specificity of the identified signatures of positive selection. There were no signatures of selection for all surface residues, suggesting that positive selection occurs specifically at interface and not all surface residues. Similar to the surface analysis, when coding variants were stratified based on the hydropathicity and polarity of the residues encoded by them, colocalization of high F_ST_ was seen only for IAV for the hydropathicity based stratification (Figures 4E and 4F) and for none of the viruses for the polarity-based metric (Supplementary Figure 3).

**Figure 4.**
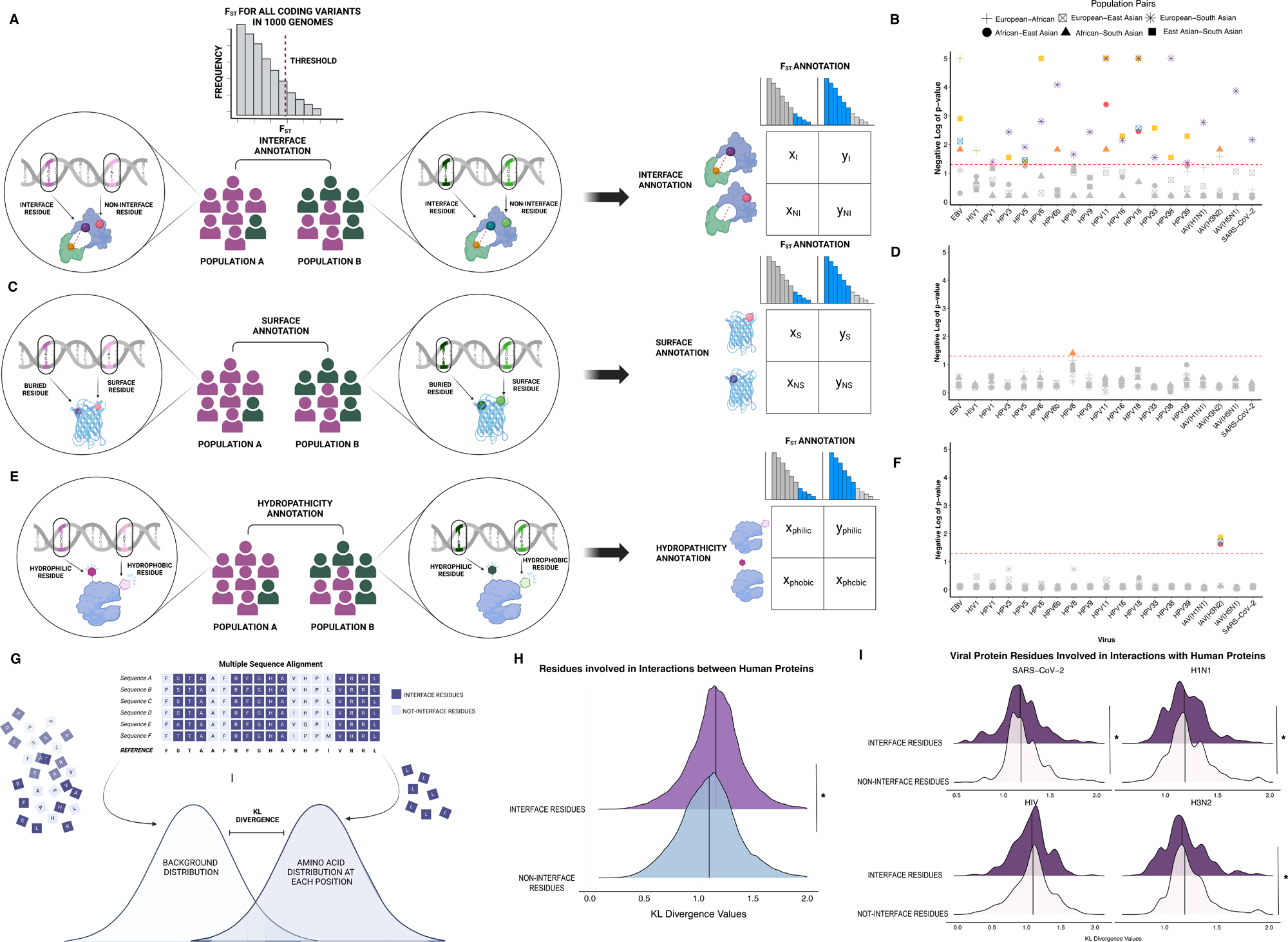
Interface Residues on Viral and Human Proteins have Differing Patterns of Selective Pressure and Conservation Compared to Non-interface Residues. (A) Schematic showing how the difference in proportion of high F_ST_ variants coding for interface and non-interface residues on human proteins was estimated. (B) Residues at interfaces of human-viral protein interactions were found to be under significantly higher selective pressure compared to not-interface residues. (C) Schematic showing how the difference in proportion of high F_ST_ variants coding for surface and buried residues on human proteins was estimated. (D) Residues at the surface of human proteins interacting with viral proteins did not have significant differences in selective pressure compared to buried residues. (E) Schematic showing how the difference in proportion of high F_ST_ variants coding for hydrophobic and hydrophilic residues on human proteins was estimated. (F) Hydrophilic residues on human proteins interacting with viral proteins did not have significant differences in selective pressure compared to hydrophobic residues. (G) Conservation for protein residues was estimated using KL Divergence. (H) Interface residues for human-human protein interactions were significantly more conserved than non-interface residues. (I) Different patterns of conservation were different for viral interface residues. * represent p-value < 0.05 for a Mann-Whitney U Test.

### Evolutionary Patterns for Interface Residues on Viral Proteins Differ from that Seen for Human Proteins

Humans and viruses are engaged in an evolutionary “arms race.” Interactions between viral and human proteins are critical for viruses but cause key alterations to host molecular phenotypes that are directly linked to viral pathogenesis. Therefore, while the human genome evolves to resist viruses, viruses are under pressure to continuously evolve to evade or modulate host immunity (Sironi et al. 2015).

We aimed to characterize the evolutionary patterns of viral protein residues using longitudinal data and compared the mode of evolution between residues at and away from the interface of protein interactions with human proteins. To this end, we used the predicted structural information from our database to compare the interface and non-interface residues for HIV1 (taxon id: 211044), SARS-CoV-2 USA-WA1/2020 (taxon id: 2697049), IAV(H1N1, taxon id: 211044), and IAV(H3N2, taxon id: 385599), given the constraints of data availability. Frequency distribution of amino acids at each position of a protein was compared to a null distribution using Kullback-Leibler divergence (Figure 4G). Positions with high divergence from random are conserved, while those showing small divergences are not conserved. When human protein residues were stratified based on their location within or away from interface regions of human protein-protein interactions, interface residues were significantly more conserved than non-interface residues (Figure 4H), which is in line with expectations as these residues play important roles in maintaining these key homeostatic interactions (Wang et al. 2012; Sahni et al. 2015). However, when the residues on viral proteins were analyzed, no significant difference in degree of conservation was observed for interface and non-interface residues for the proteins of the sexually transmitted blood borne and well established in humans HIV1 (Figure 4I). Conversely, for respiratory tract viruses emerging into the human population, non-interface residues showed higher degree of conservation compared to interface residues (Figure 4I). Viral protein residues at interaction interfaces with human proteins are less conserved compared to non-interface residues, which differs from the pattern observed for human protein residues interacting with viral proteins. These residues may well be involved in interactions with proteins of different hosts and therefore are not optimized for interacting with those of any particular one.

### Extent of Interface Sharing Differs Based on Mutation Rate

As obligate parasites, viruses have evolved various strategies to hijack host cellular machinery to facilitate their replication, and forming interactions with the host protein play an important role in this process (Lasso, Honig, and Shapira 2021). Viruses frequently imitate different structural and functional properties of host molecules to hijack or interfere with cellular processes, such as nucleic acid metabolism and immune response regulation (Lasso, Honig, and Shapira 2021). Molecular mimicry can occur at sequence or structure level, including domain and interface mimicry, with interface mimicry being the most common (Guven-Maiorov et al. 2020; Franzosa and Xia 2011; Mihalič et al. 2023). Structural mimicry is a widely used strategy by viruses, irrespective of their genome size and mode of replication, and significantly shapes host range as reported by Lasso et al who studied global structural mimicry, independent of protein interactomes, to understand how the human proteome dictates the viral structural space (Lasso, Honig, and Shapira 2021). However, this approach does not account for how viruses with diverse global structures can exploit local structural similarities to replicate human protein-protein interactions through interface mimicry. A systematic investigation of viral interface mimics is crucial for uncovering how viruses disrupt host interactomes and manipulate signaling pathways to their advantage. Since viruses within the same family often leverage conserved cellular pathways for replication through sometimes preserved host protein interactions, a systems level understanding of the conserved target protein interaction motifs can help identify shared vulnerabilities that can be targeted with antiviral therapeutics (Gordon et al. 2020).

Given the implications of shared and unique host-viral interfaces, we focused on quantifying the extent of sharing between viral-human and human-human protein interactions as well as different viral-human protein interactions across protein families using the Dice index (Figure 5A). When viruses were separated based on their observed mutation rates, we found that viruses with mutation rate >=1e^-04^ substitution/site/year had significantly higher extent of interaction mimicry compared to <1e^-04^ substitution/site/year (Figure 5B) (Holmes 2003). Counterintuitively, the viruses with higher mutation rate had higher extent of sharedness of interaction interfaces amongst themselves compared to viruses with lower mutation rates (Figure 5C). The lowest extent of shared interfaces was across viruses with higher compared to those with lower mutation rates (Figure 5C). To ensure that these patterns were not an artifact of the underlying differences in quality of predictions, we compared the distribution of the AlphaFold scores between these classes of viruses and found non-significant differences (Supplementary Figures 4A and 4B). These trends were also observed when two viruses with different mutation rates which produce slow progressing, but major disease were compared (Figures 5D and 5E). When proteins from HPVs with high cervical carcinoma risk were compared with proteins from HIV, which has a higher mutation rate, HIV proteins had a significantly higher rate of interaction mimicry (Figure 5D). However, the difference in extent of sharing amongst HIV proteins and high-risk HPV proteins was not significant, but HIV proteins tended to have a higher extent of sharedness (Figure 5E). These findings suggest that viruses with high rate of mutation not only have higher rates of interface mimicry but also bind to the same human protein interaction interfaces as their other viral counterparts (Figure 5F). Given that viruses with lower mutation rates primarily correspond to DNA viruses and those with higher mutation rates to RNA viruses, our findings are in line with prior work (Lasso, Honig, and Shapira 2021). Taken together, our results show that viruses with lower mutation rates, which often have large genomes preserve interactions by global mimicry, not necessarily interface mimicry, limiting the promiscuity of their interactions. However, viruses with higher mutation rates are better adapted to bind multiple interaction interfaces and thus fulfill the same functions with a minimalistic set of proteins.

**Figure 5.**
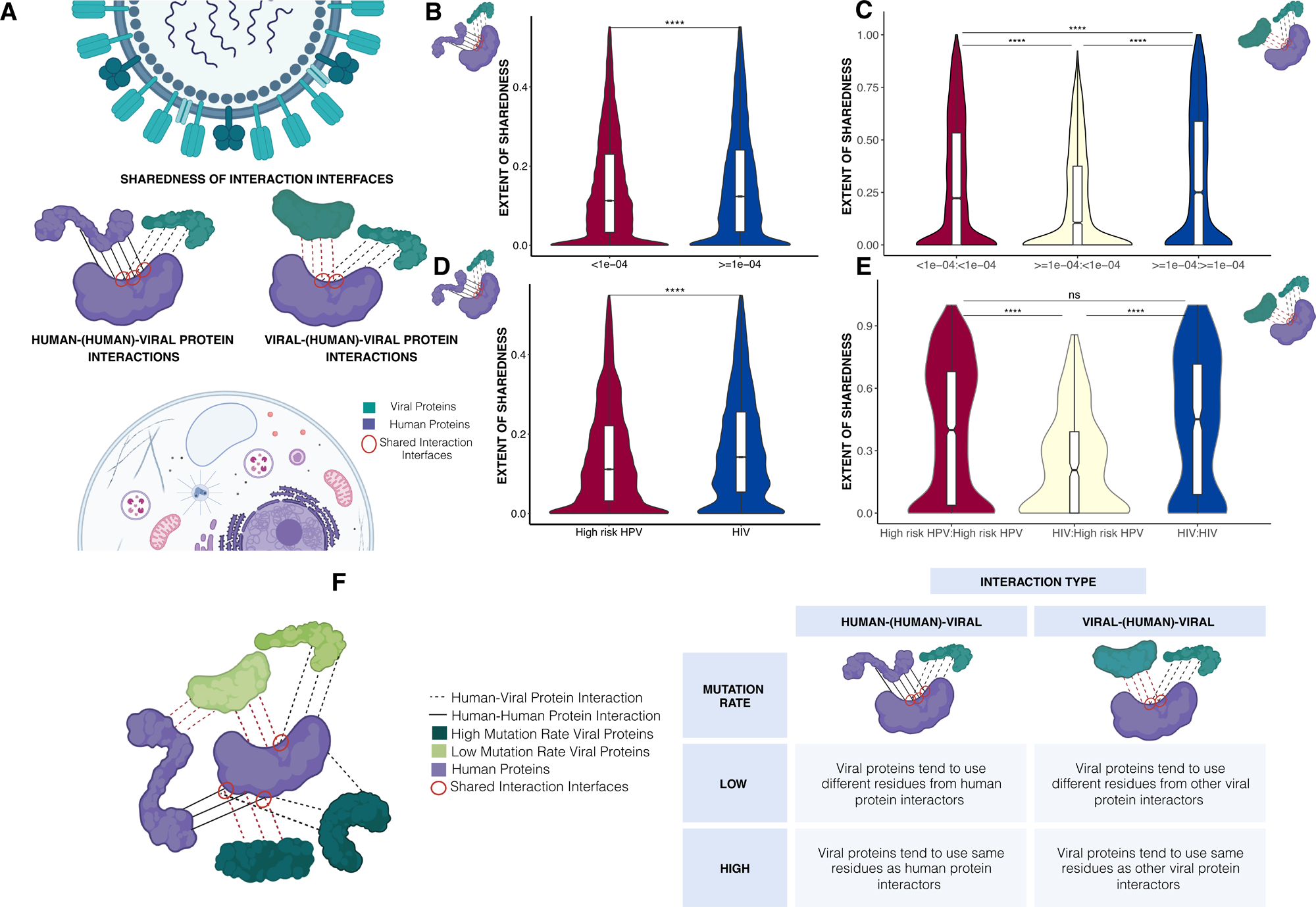
Studying the Extent of Interface Sharedness between Interactions involving Viral and Human Proteins. (A) Extent of interface sharedness was estimated for viral-human and human-human protein interactions (interface mimicry) and for different viral-human protein interactions using Dice index. Comparisons of viruses with high and low mutation rates (>=1e^-04^ and <1e^-04^ substitution/site/year, respectively) for (B) extent of overlap between human and viral proteins and (C) viral proteins binding to the same human protein are shown. Comparisons of HPV and HIV1 for (D) extent of overlap between human and viral proteins and (E) viral proteins binding to the same human protein are shown. (F) Schematic outlining the pattern of interface sharing for viruses with high and low rates of mutation. *** represent p-value < 0.05 for a Mann-Whitney U Test, adjusted for multiple comparisons where needed.

### E6 Binding Patterns Segregate High-risk and Low-risk HPVs

HPVs represent a large family of small dsDNA viruses that infect basal epithelial cells, leading to the development of benign and malignant lesions in the skin and mucosa, including those of the anogenital, upper respiratory, and digestive tracts (de Martel et al. 2017). HPVs that infect the genital tract are categorized into high and low-risk based on their oncogenicity. High-risk HPVs, such as HPV16 and HPV18, are the leading causes of cervical cancer (Longworth and Laimins 2004; Cohen et al. 2019), whereas infections by low-risk HPVs are primarily associated only with benign conditions like warts or hyperplasias, and only occasionally cause cancer (Doorbar et al. 2012). Previous studies have primarily focused on segregating pathogenic HPVs based on the sequence and expression of the oncogenes that code for E6 and E7 proteins (Doorbar et al. 2012; Schiffman et al. 2016). In their study, Lasso et al. showed that high and low-risk HPVs can be clustered based on the interactions between the viral and human proteins (Lasso et al. 2019). While this comparison of the interactomes can reveal differences in global circuits underlying disease pathogenesis, they lack the structural information that is critical for deriving mechanistic insights. Studying the interaction interfaces of these protein interactions is crucial, as they determine binding specificity, interaction strength, and the potential for competitive or allosteric regulation, providing deeper insight into the differences in the oncogenic potential of HPVs.

E6 is necessary for HPV replication and oncogenesis. During HPV infection, E6 deregulates cell cycle regulatory pathways to modify the cellular environment of terminally differentiated cells to facilitate HPV replication by binding to tumor suppressor proteins, cyclins, and cyclin-dependent kinases (Miranda Thomas et al. 2002; Mantovani and Banks 2001; Yim and Park 2005; Burd 2003; Syrjänen and Syrjänen 1999). This activity also favors carcinogenesis. Structural and functional differences between E6 from high and low-risk HPVs have been shown to contribute to the increased oncogenic potential of the former (Oh, Longworth, and Laimins 2004; Pal and Kundu 2019; Underbrink et al. 2016). Given the large number of reported E6 interactions in literature, our dataset provided us with a unique opportunity to systematically study the structural differences in the binding patterns of high and low-risk HPV E6 proteins.

We analyzed all E6 protein interactions from 23 HPVs of differing oncogenic potential, annotated based on the Centers for Disease Control and Prevention standards (Saraiya et al. 2015), to investigate how E6 proteins from different HPVs utilize human interaction interfaces. We observed a significantly lower extent of sharing between high and low risk HPVs compared to that within each group (Figure 6A), which is similar to the pattern of sharing observed across all HPV viral proteins (Supplementary Figure 5). This difference suggests that E6 from oncogenic and non-oncogenic HPVs bind to distinct domains on the same human protein, which may result in the differential downstream effects of these interactions between high and low-risk HPVs, despite all their E6 binding to the same proteins (M. Thomas and Banks 1999; Drews, Case, and Vande Pol 2019). From our list of E6 interactors, we prioritized 11 functionally important human proteins that interact with E6 from HPVs with known oncogenic classifications and quantified the association between E6 binding patterns and HPV oncogenicity using an adjusted normalized mutual information (NMI) score (Methods, Figure 6B).

**Figure 6.**
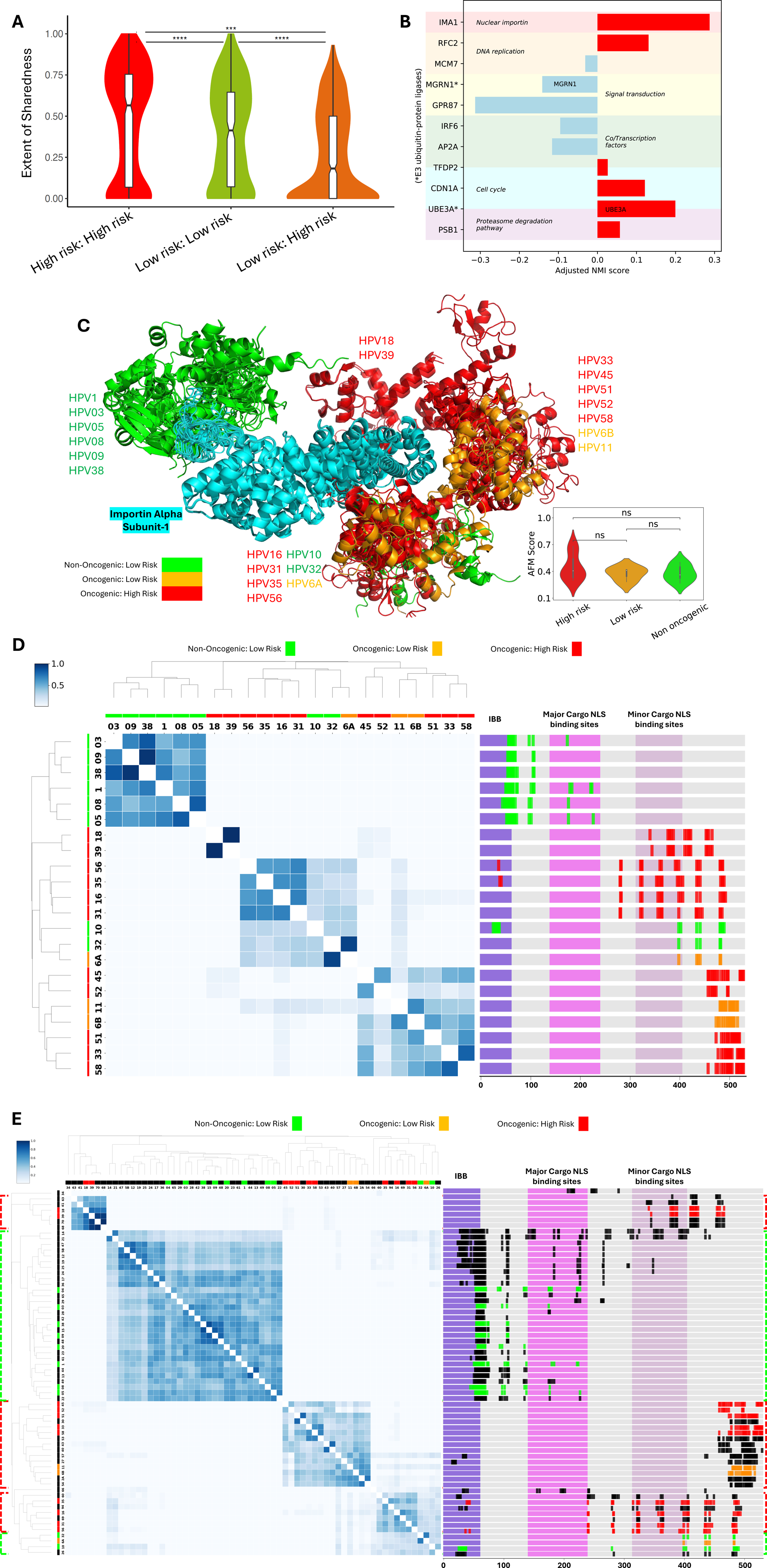
Segregation of High-Risk and Low-Risk HPVs Based on the Binding Sites of Human Proteins for Viral Protein E6. (A) Comparison of extent of interface sharing, measured by Dice index, by E6 proteins from high-risk and low-risk HPVs. (B) Adjusted NMI scores of functionally important human proteins that have intensive interactions with the E6 protein of HPVs. The functions of proteins are labelled with background colors. The adjusted NMI score measures how likely the binding sites of human proteins for E6 are consistent with the oncogenic levels (high-risk or low-risk) of HPVs. (C) Predicted structures of interactions between IMA1 and E6 of different HPVs with known oncogenic annotations are superimposed, showing groups of binding sites. HPV6 (6A and 6B) and HPV11, considered low-risk as they do not cause cervical cancer, are colored blue as they may cause other types of cancer. The right-bottom panel shows the distribution of prediction scores among different HPV groups. (D) Heatmap with hierarchical clustering plot (left panel) based on the Jaccard index of binding sites on IMA1 (right panel). The axis labels of the heatmap are the type numbers of HPVs, with their oncogenic levels indicated in red bars (high-risk) and green bars (low-risk) above the axis labels. The right panel shows the locations of binding sites of different E6 proteins on IMA1 sequences, where three key domains are annotated: IBB (importin beta binding domain, position: 2-66), NLS binding site (major, position: 142-238), and NLS binding site (minor, position: 315-403). (E) Extended heatmap with hierarchical clustering plot based on the Jaccard index of binding sites of all HPVs with known E6 sequences on IMA1. The oncogenic levels of the clusters are annotated with colored dashed rectangles across the heatmap and domain view of IMA1 (red for high-risk and green for low-risk).

Among these 11 proteins, KPNA2 had the highest adjusted NMI score, indicating a strong association between E6 binding patterns and the oncogenicity of the corresponding HPV. KPNA2 is a nuclear transport protein which consequently plays roles in cell cycle regulation, DNA repair, and transcriptional control (Alshareeda et al. 2015; Christiansen and Dyrskjøt 2013; Ma and Zhao 2014). Overexpression of *KPNA2* has been implicated in cervical cancer development and progression via deregulation of the E2F/Rb pathway (van der Watt, Ngarande, and Leaner 2011). When the co-complex structures of binary interactions between KPNA2 and E6 from 23 HPV were superimposed based on the structure of KPNA2 (cyan, Figure 6C), E6 proteins binding to KPNA2 could be clustered into four groups: one group of non-oncogenic HPVs, one of oncogenic high-risk HPVs, one of oncogenic low and high-risk HPV, and one of non-oncogenic and oncogenic low and high-risk HPVs. To ensure that this clustering was not influenced by structural prediction biases, we compared the distributions of AlphaFold structure quality scores across different HPV categories and found no significant differences (right-bottom of Figure 6C), confirming that the clustering patterns reflect biologically relevant variations in E6-KPNA2 binding interfaces. The individual co-complex structures are presented in Supplementary Figure 6.

To better understand this clustering, we calculated the Jaccard Index for E6 binding sites on KPNA2 across different HPV strains and visualized the results as a heatmap with a hierarchical clustering dendrogram (left panel of Figure 6D). The two mixed groups comprising HPV strains of differing oncogenic potential could be further subdivided into smaller clusters of similar oncogenic potential along the dendrogram. Visualization of the interface residues on the KPNA2 sequence (right panel of Figure 6D) revealed that none of the high-risk HPV binds to the N-terminal region that binds to importin beta (IBB domain). Instead, they interact with the minor cargo nuclear localization signal (NLS) binding domain. In contrast, E6 proteins from non-oncogenic HPVs preferentially bind to the IBB. Given that nuclear import is crucial for oncogenic functions of E6, the preferential binding of benign E6 proteins to the IBB, potentially interrupting the importin-α/importin-β interaction required for nuclear import, may play a role in their reduced oncogenic potential (Pal and Kundu 2019; Le Roux and Moroianu 2003; Yi et al. 2020). Our findings align with those of Mespiede et al. (Mesplède et al. 2012) who showed that E6 from high-risk HPVs predominantly localized in the nucleus, while benign E6 proteins were found in both the cytoplasm and nucleus.

These results indicate that the co-complex structure of the interactions between E6 and KPNA2 and the binding patterns of E6 on KPNA2 provide strong predictive power for the pathogenicity of HPVs. We repeated the analysis and expanded the scope of the analysis to 64 HPVs with E6 sequences in UniProtKB/Swiss-Prot. We calculated the Jaccard Index for all 64 E6 proteins based on the binding sites on KPNA2 to plot a heatmap with a hierarchical clustering dendrogram (Figure 6E). We identified five clusters based on this heatmap, which were labeled by the HPVs with known pathogenic annotations using a majority voting strategy. In this way, we predicted the pathogenic level for all HPVs included in the analyses. For example, HPV41 and HPV63 are likely high-risk oncogenic (consistently with the original suggestion by Zur Hausen for HPV41(Grimmel et al. 1988)), and HPV23, HPV24, and HPV28 are likely non-oncogenic. The list of predicted oncogenic levels of all HPVs is provided in Supplementary Table 1.

### HSV-1 UL37 competes with MAVS for TRAF6 binding to inhibit IFN activation

Our analysis identified interactions between the UL37 inner tegument protein of HSV-1 and the human protein MAVS on the same interface on MAVS that interacts with TRAF6. TRAF6 and MAVS are components of the RIG-I-like receptor (RLR) signaling pathway which detects viral RNA to activate antiviral innate immunity (Figures 7A-B). Detection of viral RNA by RIG-I-like receptors triggers the formation of a signaling platform centered on the adaptor MAVS. MAVS recruits the E3-ubiquitin ligase TRAF6 through a consensus TRAF6-binding motif within its central disordered region. TRAF6 engagement at the MAVS signalosome is required for the activation of the transcription factors IRF3 and NF-kB which induce interferons (IFNs), key cytokines that establish an antiviral state (Xu et al. 2005; S. Liu et al. 2013; Fang et al. 2017). During HSV-1 infection, UL37 engages TRAF6 through a TRAF6-binding motif in its disordered C-terminal tail to activate NF-κB, which is required for viral replication (Xueqiao Liu et al. 2008). Additionally, UL37 inhibits IFN induction by deamidating and inactivating the sensors RIG-I and cGAS during HSV-1 infection (Zhao et al. 2016; Zhang et al. 2018).

**Figure 7.**
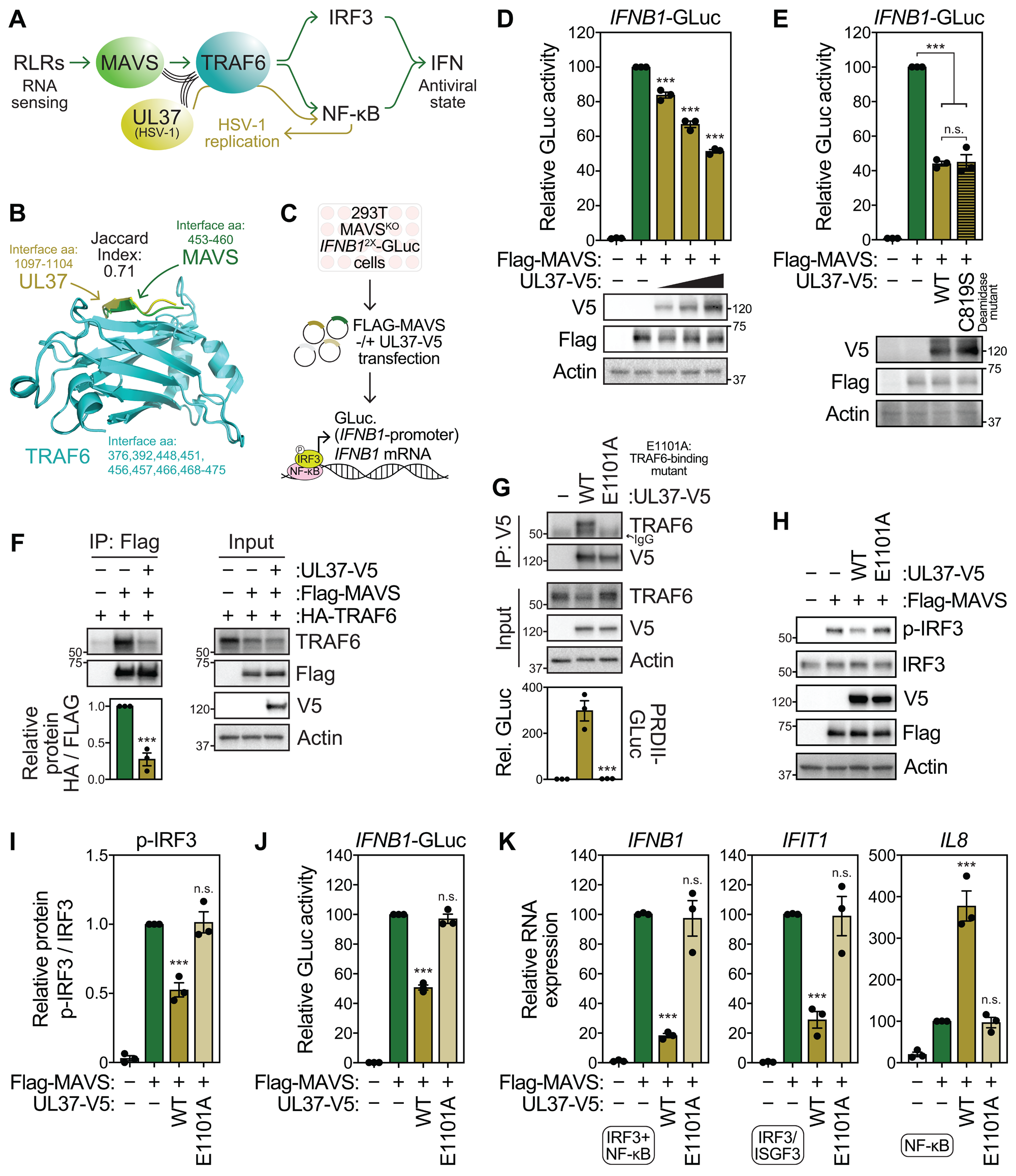
HSV-1 UL37 competes with MAVS for TRAF6 binding to inhibit IFN activation. (A) Schematic of RIG-I-like receptor and Herpes simplex virus-1 UL37 signaling. (B) Model demonstrating shared interface residues on TRAF6 for UL37 (predicted by Alphafold) and MAVS (PDB: 4Z8M) binding. (C) Schematic of experiments investigating UL37 suppression of MAVS signaling. 293T MAVSKO IFNB1-GLuc reporter cells induce Gaussia luciferase (GLuc) driven by tandem IFNB1 promoter elements after MAVS overexpression. (D) (Top) Relative GLuc in the supernatant (S/N) from 293T MAVSKO IFNB1-GLuc cells following overexpression of Flag-MAVS and increasing doses of UL37-V5 at 36 hours post-transfection (hpt). (Bottom) Immunoblot analysis of lysates from cells treated in the indicated manner. (E) (Top) Relative GLuc in the S/N from 293T MAVSKO IFNB1-GLuc cells following overexpression of Flag-MAVS and UL37-V5 WT or C819S deamidation mutant at 36 hpt. (Bottom) Immunoblot analysis of lysates from cells treated in the indicated manner. (F) Immunoblot analysis of anti-Flag and anti-HA immunoprecipitated extracts as well as input lysates from 293T MAVSKO cells transfected with HA-TRAF6, Flag-MAVS, and UL37-V5 as indicated (24 hpt). Quantification of HA-TRAF6 relative to immunoprecipitated Flag-MAVS and Flag-MAVS relative to immunoprecipitated HA-TRAF6 are shown below as indicated. (G) (Top) Immunoblot analysis of anti-V5 immunoprecipitated extracts and input lysates from 293T PRDII-GLuc (NF-κB reporter) cells expressing UL37-V5 WT or E1101A TRAF6-binding mutant (24 hpt). (Bottom) Relative GLuc in S/N from 293T PRDII-GLuc cells expressing UL37-V5 WT or E1101A. (H) Immunoblot analysis of 293T MAVSKO IFNB1-GLuc cells following expression of Flag-MAVS as well as UL37-V5 WT and E1101A at 36 hpt. (I) Quantification of phosphorylated IRF3 (S386) relative to total IRF3 from experiments in (H). (J) Relative GLuc in the S/N from 293T MAVSKO IFNB1-GLuc cells following overexpression of Flag-MAVS and UL37-V5 WT and E1101A (36 hpt). (K) RT-qPCR analysis of IFNB1 (target of IRF3 and NF-κB), IFIT1 (target of IRF3 and ISGF3 after activation by interferons), and IL8 (target of NF-κB) mRNA expression relative to HPRT1 from 293T MAVSKO IFNB1-GLuc cells following overexpression of Flag-MAVS and UL37-V5 WT and E1101A (36 hpt).

We hypothesized that UL37 could further interfere with the RLR-MAVS pathway by competing with MAVS for TRAF6 binding to limit IRF3 activation and thus IFN induction. To determine whether UL37 can inhibit signaling through MAVS independently of upstream modulation of RIG-I, we tested whether UL37 can alter IFN induction after MAVS overexpression in 293T MAVSKO *IFNB1*-GLuc reporter cells (Figure 7C). MAVS overexpression is sufficient to drive signaling independently of RIG-I activation and *Gaussia* luciferase (GLuc) induction in these cells is under the control of *IFNB1* promoter elements (Gokhale et al. 2024). Overexpression of UL37 inhibited GLuc induction after MAVS overexpression in a dose dependent manner (Figure 7D). Both wild-type (WT) and deamidase mutant (C819S) UL37 inhibited antiviral signaling equally, which indicates that the suppression of MAVS signaling by UL37 was independent of its enzymatic activity (Figure 7E). Indeed, co-immunoprecipitation analysis revealed that UL37 inhibits the association between MAVS and TRAF6 (Figure 7F).

To test whether TRAF6-binding by UL37 was critical for the suppression of MAVS signaling, we generated a TRAF6-binding mutant UL37 (E1101A) that neither bound TRAF6 nor activated NF-κB (Fig. 7G) (Xueqiao Liu et al. 2008). While overexpression of WT UL37 suppressed the phosphorylation of IRF3 at S386, a marker of IRF3 activation upon MAVS signaling, overexpression of UL37 E1101A did not (Fig. 7F). Similarly, UL37 E1101A did not inhibit the induction of *IFNB1*-GLuc or *IFNB1* and *IFIT1* transcripts which require IRF3 activation unlike WT UL37 (Fig. 7G-H). Moreover, UL37 WT but not UL37 E1101A enhanced the expression of the NF-kB-target gene *IL8* (Fig. 7H). Together, these data indicate that HSV-1 UL37 limits the association between TRAF6 and the MAVS signalosome, thereby inhibiting the activation of IRF3 and consequently, IFN production. This effect likely contributes to the inability of HSV-1 lacking UL37 to inhibit IFN induction during infection (Zhao et al. 2016; Zhang et al. 2018).

## Discussion

Interactions between viral and human proteins are required for viral replication and pathogenesis. Analyzing the structures of these interactions provides key insights into the mechanisms utilized by viruses to trigger and evade host defenses (de Chassey et al. 2014; Cakir et al. 2021; Wierbowski et al. 2021). We collected a comprehensive set of 11,666 binary physical viral-human and 2,838 viral-viral protein-protein interactions for all human-infecting viruses in the literature. However, approximately 95% of these protein interactions lack structural information. After comparing multiple state-of-the-art methods, we applied AlphaFold (Jumper et al. 2021; Abramson et al. 2024; Evans et al. 2022) to model all these interactions to create the most comprehensive 3D viral interactomes till date (Jumper et al. 2021; Abramson et al. 2024; Evans et al. 2022; Lin et al. 2023; Ketata et al. 2023; Dominguez, Boelens, and Bonvin 2003). Analysis of this large-scale dataset revealed unique evolutionary features of the host and viral interaction interfaces, distinct patterns of interface mimicry by viral proteins based on mutation rates, and characteristic differences in interaction domains between oncogenic and benign HPVs. Furthermore, we uncovered a novel mechanism through which UL37 disrupts the innate immune response.

The evolutionary ‘arms race’ between humans and viruses results in a continuous cycle of adaptation and counter-adaptation shaping the genomes of both. Infections by pathogens, including viruses, exert selective pressure on the human genome driving population divergence (Van Blerkom 2003). Several studies have shown higher rates of selective pressure on immune genes, particularly those that encode proteins that directly interact with viral proteins (Meyerson et al. 2014; Barreiro et al. 2009; Lasso et al. 2019). Exome scans comparing single-locus estimates of FST with the exome-wide background revealed higher rates of selection at loci coding for interface residues of human-viral protein interactions compared to other residues, reflecting their functional importance in mediating these interactions and modulating the downstream signalling cascades (Chhibbar et al. 2024). However, viral protein residues were found to have lower or not significantly different conservation rates for interface residues compared to non-interface residues—indicating the need for rapid adaptation at these sites to evade or modulate host immunity as well as to replicate in different species.

One common mechanism adopted by viral proteins to modulate host immune response and improve fitness is molecular mimicry. Structural mimicry is a common strategy employed by viruses, regardless of their genome size or replication method, and plays a crucial role in determining host range as reported by Lasso et al. (Lasso et al. 2019). A systematic investigation of the patterns of interface sharing between viral-human and human-human as well as different viral-human protein interactions revealed that viruses with high mutation rates ensure higher interaction promiscuity by exploring different interaction interfaces than other viral proteins and have higher rates of interface mimicry compared to viruses with lower mutation rates. Differences in patterns of structural mimicry was also observed for HPVs based on oncogenic risk. Quantifying the relationship between the binding patterns of the oncoprotein E6 from different HPVs and their oncogenic potential revealed a strong association for several functionally important proteins. In particular, the interaction with KPNA2 revealed distinct interface usage patterns segregating high– and low-risk HPVs, underscoring the role of nuclear import in oncogenesis of HPV. By systematically clustering HPVs based on E6 binding interfaces on multiple human proteins, we predict oncogenic risk for uncharacterized HPVs. These differential surface interactions highlight how different HPVs of varying oncogenic risk differ not only in global patterns of protein interactions but also in the specific interface usage (Lasso et al. 2019), and it can augment sequence-based approaches for pathogen classification and risk assessment, for HPV as well as other viruses.

While systemic analysis of the dataset revealed key insights into pan-viral mechanisms of pathogenesis, targeted examination of specific interactions allowed us to generate testable mechanistic hypotheses for individual viral proteins. Analyzing the structure of the interaction between UL37, a HSV-1 protein, and TRAF6, which is critical for HSV-1 replication, revealed significant mimicry of the MAVS binding region on TRAF6. TRAF6 and MAVS interaction is critical for induction of IFNs to establish an antiviral state (Xu et al. 2005; S. Liu et al. 2013; Fang et al. 2017). Using mutant and WT UL37, we showed that TRAF6 binding by UL37 suppresses MAVS signaling and, subsequently, IFN activation. The inhibition of IFNs by the deamidation activity of UL37 has been previously reported, but we show a novel mechanism through which UL37 inhibits IFNs (Zhao et al. 2016; Zhang et al. 2018).

Through systematic and targeted analysis of our 3D interactome, we demonstrated that this new resource can significantly expand the scope of viral research, facilitating mechanistic interpretation and experimental validation of critical hotspots that are likely to influence viral replication, immune evasion, or disease outcomes. Despite its great potential, limitations still remain, presenting opportunities for future improvements. Evaluating predictions for multi-subunit viral-human complexes is challenging. Therefore, our dataset is currently limited to binary interactions. However, integrating additional data sources, such as the structurally resolved human protein interactome, can potentially overcome this limitation and enable the analysis of multi-component assemblies, uncovering cooperative or competitive interactions that influence infection dynamics. Moreover, while computational tools, such as AlphaFold, can be powerful tools to address limitations of experimental methods, biochemical and biophysical assays are essential for refining or validating these models, particularly when predicted structures suggest novel interaction modes. Nevertheless, through extensive benchmarking we have shown that AlphaFold is currently the best tool for accurate structure prediction, showing high concordance between predicted and real structures of the benchmark set of interactions. We and others are actively developing new computational tools that improve current interface prediction approaches (Xiong et al. 2024). These tools can readily be incorporated into our 3D viral interactome modeling pipeline to enhance downstream inference.

We present the largest and most comprehensive resource of 3D structural models for host-viral protein interactions till date. Through systematic analysis of this structural data, we uncovered unique functional and evolutionary features of critical interfaces, providing novel insights into the molecular mechanisms underlying viral infection and pathogenesis. Our collection of interactomes offers a unique platform for integrating multi-omic data to investigate how genetic and proteomic variations in host immune factors can impact viral interactions and modulate disease response. This integrated approach can help researchers uncover crucial pan-viral interaction patterns and formulate testable hypotheses, ultimately contributing to the development of new therapeutic targets and treatment strategies.

## Methods

### Data collection

#### Benchmark datasets

We used the list of known pathogen-host interactions from one of our previous works (Wierbowski et al. 2021), containing 509 viral-human and bacteria-human interactions with known co-complex structures in the PDB database. To facilitate benchmarking of docking and docking-like methods, we also prepared structures of individual proteins that have the highest sequence coverage and are not part of any co-complex structures. For intra-species interactions, we randomly selected 200 human-human interactions with available PDB structures.

### 3D structure prediction for binary interaction

#### AlphaFold-Multimer

We utilized AlphaFold-Multimer v2.2.0, downloaded on March 10, 2022, along with all requisite databases from the AlphaFold GitHub (https://github.com/google-deepmind/alphafold). Installation was performed on our local machines. Structure predictions were executed using the pre-trained models (trained on all PDBs released before April 2018) with default parameters.

#### ESMFold

For predicting the 3D structures of binary interactions using ESMFold, we concatenated the sequences of individual proteins using a colon “:” separator, indicating a significant gap and guiding the model to output two distinct chains. ESMFold was downloaded from the ESM GitHub repository (https://github.com/facebookresearch/esm) on March 20, 2023, and predictions were made using the default settings of the trained model.

#### DiffDock-PP

We obtained DiffDock-PP from its GitHub repository (https://github.com/ketatam/DiffDock-PP) on January 24, 2024, and followed the provided installation instructions. The co-complex structures were predicted using the trained models with default parameters, utilizing the prepared single protein structures from the benchmark dataset.

#### HADDOCK and ECLAIR

Docking simulations were performed using HADDOCK 2.4 (release date: September 2020). We conducted both unguided docking and docking guided by predicted interface residues. For the latter, interface residues were predicted using our previously developed method, ECLAIR (Meyer et al. 2018).

### Interface prediction

For methods generating complete co-complex structures, we identified interface residues by evaluating their solvent-accessible surface area (SASA). Residues were classified as interface residues if their SASA exceeded 15 square angstroms and the difference in SASA between their bound and unbound states was greater than 1 square angstrom.

For interface prediction using DELPHI, D-Script, and DLPred, we downloaded the respective packages from their sources: DELPHI from GitHub (https://github.com/lucian-ilie/DELPHI), D-Script from GitHub (https://github.com/samsledje/D-SCRIPT), and DLPred from their website (http://qianglab.scst.suda.edu.cn/dlp/). Predictions were conducted using default parameters for all methods. As there is no universal cutoff for prediction scores, we selected the cutoff yielding the highest F1 score in our benchmark for each method.

### Measurement metrics

To comprehensively evaluate the structure-level qualities of the predicted co-complex structures, we employed a suite of metrics, including DockQ score, TM-Score, RMSD, iRMSD (interface RMSD), and lRMSD (ligand RMSD). TM-Score and RMSD were calculated using the open-source tool MM-align (Mukherjee and Zhang 2009), while DockQ, iRMSD, and lRMSD were computed using the DockQ package (Basu and Wallner 2016). For the predicted interface residues, we calculated individual precisions and recalls for each interaction. Additionally, we computed the overall precision and recall for all interactions combined, representing the macro average of the individual precisions and recalls.

To fairly evaluate the segregation of binding sites on the same human protein by E6 proteins from high-risk and low-risk HPVs—particularly when the number of segregated clusters differs—we adjusted the normalized mutual information (NMI) by dividing it by the average NMI values obtained from random permutations with the same number of clusters and obtained the so-called adjusted NMI score.

### Estimating F*_ST_* for interface and non-interface variants

#### Dataset Processing and F*_ST_* Estimation

The 1000 Genomes Phase 3 dataset was obtained from the UCSC Genome Browser (Perez et al. 2025), and samples were grouped into four superpopulations: European (EUR), South Asian (SAS), East Asian (EAS), and African (AFR). Coding variants were annotated using BISQUE (Meyer, Geske, and Yu 2016). Pairwise F_”#_ values for each variant were computed using VCFtools (v0.1.15) (Danecek et al. 2011) for the following population comparisons: EUR vs. SAS, SAS vs. EAS, EAS vs. AFR, and AFR vs. EUR. The 75th percentile F_”#_ threshold was then determined.

#### Selective pressure analysis on viral interaction interfaces

For viruses with at least 20 predictions, coding region variants in human interactors were classified as interface or non-interface. To assess selective pressure on viral interaction interfaces, we compared the proportion of variants in interface and non-interface regions with F_”#_ values above the 75th percentile across population pairs.

### Estimating KL divergence values for interface and non-interface variants

#### Sequence retrieval and alignment

Protein sequences for selected viruses (H1N1, H3N2, HIV, and SARS-CoV-2) were downloaded from the NCBI Virus database (“NCBI Virus,” n.d.). The taxonomy ID with the highest number of predictions in our database was used for sequence selection. These sequences were aligned to their respective reference proteins (used for interface prediction) using MAFFT (v7.471) (Katoh et al. 2002).

#### Residue Classification and KL Divergence Calculation

Aligned residues were classified as interface or non-interface based on our predictions. KL divergence was computed at each position to measure how amino acid distributions deviate from the background distribution. The KL divergence, *D_KL_*, is defined as:

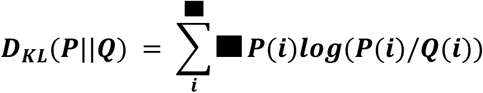

where P(*i*) is the frequency of amino acid *i* at a given position, and *Q*(*i*) is its frequency in the background distribution.

#### Statistical Comparison

We used the Mann-Whitney U test to compare KL divergence between interface and non-interface residues, assessing differences in evolutionary constraint across these regions.

### Virus Mutation Rate Classification

To evaluate the impact of viral mutation rates on interface conservation, viruses were categorized based on nucleotide substitution rates per site per year, extracted from peer-reviewed studies (Supplementary Table 2). For consistency and interpretability, mutation rates were rounded to representative magnitudes, rather than using their precise values. Viruses were grouped into two categories: < 1 *e*^-4^ and >= 1 *e*^-4^ based on the order of magnitude of the mutation rate. This classification allowed us to analyze whether rapidly evolving viruses exhibit lower interface conservation compared to more stable viruses.

### Estimating Extent of Interface Sharing between PPI Pairs

To quantify interface residue overlaps between interacting protein pairs, we used the Dice Index (DI):

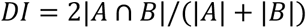

where |*A* ∩ *B*| is the number of shared interface residues, and |*A*| and |*B*| are the total interface residues in each protein.

### Experimental settings

#### Cell lines and cell culture

293T PRDII-GLuc, 293T MAVS^KO^ (Gokhale et al. 2024), and 293T MAVS^KO^ *IFNB1*-GLuc cells were grown in Dulbecco’s modified Eagle’s medium (DMEM; Thermo Fisher) supplemented with 10% fetal bovine serum (HyClone), 25 mM HEPES (Thermo Fisher), 1X non-essential amino acids (Thermo Fisher), and 1X PSG (Thermo Fisher). Cells were verified as mycoplasma free using the LookOut Mycoplasma PCR detection kit (Sigma-Aldrich). 293T MAVS^KO^ *IFNB1*-GLuc were generated by lentiviral transduction of 293T MAVS^KO^ cells as described previously (Gokhale et al. 2024).

#### Cloning and plasmids

pEFTak Flag-MAVS and Flag-empty vector (EV) were described previously (Gokhale et al. 2024). pEFTak HA-TRAF6 was generated by insertion of the TRAF6 coding sequence amplified from cDNA into *NotI*– and *PmeI*-cut pEFTak HA-EV by Infusion cloning (Takara). pEFTak UL37-V5 was generated by insertion of UL37-V5 amplified from pCDH UL37-V5 (Gift of Dr. Pingui Feng) into *KpnI*– and *PmeI*-cut pEFTak Flag-EV by Infusion cloning. pEFTak UL37-V5 C819S and E1101A were generated by site-directed mutagenesis of pEFTak UL37-V5 (WT) using the Quikchange Lightning kit (Takara). pEFTak V5-EV was generated by Infusion cloning of annealed primers containing V5 and His tag sequences into *KpnI*– and *PmeI*-cut pEFTak Flag-EV. All primer sequences are found in Supplementary Table 3.

#### Co-immunoprecipitation

For immunoprecipitation of Flag-MAVS and HA-TRAF6, 293T MAVS^KO^ cells seeded in 10 cm plates (8×10^6^ cells/plate) were transfected with 6 µg each of pEFTak HA-TRAF6 and pEFTak Flag-MAVS and 8 µg pEFTak UL37-V5 using TransIT X2 transfection reagent (Mirus) as per the manufacturer’s protocol. Total amount of DNA in each transfection was made up to 20 µg per plate with pEFTak V5-EV. At 24 hpt, cells were lysed in 500 µL IP lysis buffer (25 mM HEPES pH 7.5, 150 mM NaCl, 3 mM MgCl_2_, 2 mM EGTA, and 0.5% Triton X-100) supplemented with protease-phosphatase inhibitor cocktail (Sigma-Aldrich) for 15 mins on ice. After clarification of lysates by centrifugation at 8000 xg at 4°C, protein levels were quantified by Bradford assay (Bio-Rad). 5% of normalized lysates were kept for input samples. 500 µg of each lysate was incubated separately with anti-Flag magnetic beads (Thermo Fisher) in a final volume of 400 µL IP lysis buffer for 4 hrs at 4°C with head-over-tail rotation. Beads were washed 1X with IP lysis buffer and 3X with cold PBS. Immunoprecipitated proteins were eluted by boiling in 2X Laemmli buffer for 5 mins and subjected to immunoblotting.

For immunoprecipitation of UL37-V5, 293T PRDII-GLuc cells seeded in 6-well plates (1.6×10^6^ cells/well) were transfected with 2 µg pEFTak UL37-V5 WT or E1101A (or pEFTak V5-EV) using TransIT X2. At 24 hpt, cells were lysed in 250 µL IP lysis buffer, and protein levels were quantified after clarification. 250 µg protein lysates were incubated with Protein G Dynabeads (Thermo Fisher) pre-bound to rabbit anti-V5 antibody (Cell Signaling) for 4 hrs at 4°C with head-over-tail rotation. Beads were washed 3X with IP lysis buffer. Immunoprecipitated proteins were eluted by boiling in 2X Laemmli buffer for 5 mins and subjected to immunoblotting.

#### Gaussia luciferase reporter assays

293T MAVS^KO^ *IFNB1*-GLuc or 293T PRDII-GLuc cells seeded in24-well plates (4×10^5^ cells/well) were transfected with the indicated plasmids using the TransIT X2 reagent and media on cells was changed at 6 hpt. At 36 hpt, 10 µL supernatant from each well was transferred to white opaque 96-well plates in technical duplicate. S/N was mixed with *Gaussia* Glow assay buffer with 1X coelenterazine (Thermo Fisher), and luminescence was read on a Biotek Synergy plate reader. After removal of S/Ns for analysis, treated cells were subjected to immunoblotting and RT-qPCR analysis.

#### Immunoblotting

Whole cell lysates were prepared in a modified RIPA buffer (10 mM Tris pH 7.4, 150 mM NaCl, 0.5% sodium deoxycholate, and 1% Triton X-100) supplemented with protease-phosphatase inhibitor cocktail and clarified by centrifugation at 8000 xg for 10 mins at 4°C. Protein concentration was determined by Bradford assay and equal amounts of protein sample in 1X Laemmli buffer with 2.5% β-ME were prepared by boiling. Samples were resolved by SDS-PAGE (Tris-Glycine gels; Bio-Rad) and transferred to methanol-activated PVDF membranes (Bio-Rad) by wet transfer. Transferred membranes were incubated with relevant primary antibodies in 3% BSA in TBS-T with shaking for 1-2 hrs at room temperature or overnight at 4°C. After washing three times with TBS-T, membranes were incubated with species-specific horseradish peroxidase-conjugated secondary antibodies (Jackson, 1:5000). Chemiluminescence was detected using a Bio-Rad ChemiDoc XRS+ imaging instrument. The following primary antibodies were used for immunoblotting: rabbit anti-TRAF6 (CST, 1:1000), rabbit anti-IRF3 (CST, 1:1000), rabbit anti-IRF3 phospho-S386 (Abcam, 1:1000), HRP-conjugated anti-FLAG (Sigma-Aldrich, 1:5000), HRP-conjugated anti-V5 (Proteintech, 1:5000) and HRP-conjugated anti-β-actin (CST, 1:5000). Gels were quantified by densitometry using FIJI.

#### RT-qPCR

RNA was extracted using TRIzol (Thermo Fisher) and cDNA was generated using the PrimeScript RT-PCR kit (Takara). RT-qPCR was performed using a Viaa7 Real Time PCR instrument (Applied Biosciences) using TaqMan Universal PCR master mix II – UNG (Thermo Fisher). Primer-probe sets (IDT) are described in Supplementary Table 4.

### The 3D Viral-Human Structural Interactome Web Server

We constructed the 3D Viral-Human Structural Interactome web server, https://3d-viralhuman.yulab.org/, to provide the AlphaFold predictions as a comprehensive resource to the public. All results and raw data described herein are directly available for bulk download (https://3d-viralhuman.yulab.org/downloads). Users can quickly search specific interactions or interactomes of interest through four types of input modes:

1. **Select a prediction** from the list of all interactions or interactions within a specific virus family; go to the interaction view.
2. **Extract an interaction**, if it exists, by inputting the names/UniProt IDs of both viral and human proteins; go to the interaction view.
3. **Retrieve an interactome** by inputting the name/ID of a specific protein; go to the interactome view.
4. **Retrieve an interactome** by inputting a virus taxonomy ID; go to the interactome view.

The interactome display (Supplement Figure 7 top left and right) provides both graphical and tabular views of all involved interactions and their details. Users can interact with the graph/table by moving the cursor to specific graph nodes, edges, or table rows to show the entry names of proteins, dragging the nodes to change their positions and organizations, clicking on graph nodes to redirect to another interactome view for the protein node clicked, or clicking on graph edges or table rows to redirect to the interaction view of the specific interactions.

The single interaction view (Supplement Figure 7 bottom panel) offers comprehensive details for viral and human proteins, including structural visuals (either docked or predicted, switchable using a top-center button) and summaries of interface residues. Users can modify the list of interface residues by adjusting SASA, dSASA, and pLDDT values via sliding bars. By default, interface residues are set with thresholds of SASA ≥ 15, dSASA ≥ 1, and pLDDT ≥ 50, and are displayed in dark blue (viral) and dark green (human).

Furthermore, the interface view displays a linear sequence of both viral and human proteins for the query interaction, with interface residues on viral and human proteins marked in dark blue and dark green, respectively. It also visualizes the interfaces for other interactors of the protein below and provides Jaccard Index scores to indicate their overlaps, facilitating comparison. There are also clickable taxonomy IDs for viral proteins directing the viral interactome view, clickable UniProt IDs for both proteins directing their single protein interactome views, and clickable entry names of other interactors to direct the corresponding single interaction views.

## Supplemental Information

Supplementary Figure 1. (A) Comparison of multiple sequence alignments (MSAs) for viral-human and bacteria-human interactions. The bars display the percentage of interactions with individual and joint MSAs, while the violin plots show the distributions of MSA depths. (B) Scatter plots showing precision and recall of predicted interface residues for individual interactions that were not used in the training of AlphaFold. Contour lines indicate the density of predictions for both inter-species and intra-species interactions. (C) Comparison of structure-level quality measurements, including interface RMSD (iRMSD) and ligand RMSD (lRMSD), between interspecies and intraspecies interactions that were not used in the training of AlphaFold. (D) The comparison between AlphaFold-Multimer (AFM) and AlphaFold3 (AF3) in terms of precision (y-axis), recall (x-axis), and F1-score (contour lines) for the predicted interfaces. Smaller dots represent the performance on individual viral-human interactions, while larger dots indicate the overall performance, calculated by integrating the interfaces of all interactions and computing the corresponding precision and recall values.

**Supplementary Figure 2.** Scatter plot showing the precision and recall of predictions for the interactions with known PDB structures released after the training timestamp (April 2018) of AlphaFold.

**Supplementary Figure 3.** Difference in proportion of high F_ST_ for variants stratified by polarity of residue encoded. No significant differences were observed for any of the viruses.

Supplementary Figure 4. Distribution of AlphaFold score for interactions used in analysis of extent of sharedness between (A) viral and human and (B) viral proteins interacting with the same human proteins.

Supplementary Figure 5. Comparison of extent of interface sharing, measured by Dice index, by viral proteins from high-risk and low-risk HPVs.

**Supplementary Figure 6.** Predicted structures of the interactions between IMA1 and E6 proteins from different HPV types, categorized by their known oncogenic annotations, are grouped and presented separately. The binding domains of IMA1 involved in each interaction are highlighted in corresponding colors to indicate the specific regions engaged in binding.

**Supplementary Figure 7.** Overview of the 3D-ViralHuman Structural Interactome Browser. This figure illustrates the result pages for single interactions or interactomes in our 3D-ViralHuman structural interactome browser. The browser allows users to search for single interactions by selecting from an existing list of interactions within a specific virus family or by inputting protein names/IDs. Additionally, interactomes can be searched using a single protein or a virus taxonomy ID. The interactome display (top panels) provides both graphical and tabular views of all involved interactions and their details. Users can click on graph nodes, edges, or table rows to display specific interaction information. The single interaction display (bottom panel) includes detailed information for both viral and human proteins, showing structural displays (either docked or predicted, switchable with the button at the top middle) and summarizing interface residues for both proteins. The list of interface residues can be adjusted based on SASA value, dSASA value, and pLDDT value settings using sliding bars. By default, interface residues are defined with settings of SASA ≥ 15, dSASA ≥ 1, and pLDDT ≥ 50, and are colored dark blue (viral) and dark green (human). The interface view also presents a linear representation of the protein sequence, with interface residues on viral and human proteins annotated in dark blue and dark green, respectively. Interfaces for other interactors of the protein are displayed below for easy comparison, with the Jaccard Index indicating the overlap between interface residues of the current and other interactors. Additional features include clickable taxonomy IDs of viral proteins for the viral interactome view, UniProt IDs of both proteins for their single protein interactome views, and entry names of other interactors for the corresponding interaction views.

**Supplementary Table 1.** The oncogenic levels of all HPVs shown in the heatmap of Figure 6E

**Supplementary Table 2.** Mutation rates of viral families retrieved from literature

**Supplementary Table 3.** Primer sequences

**Supplementary Table 3.** Primer probe sets

## Resource Availability

### Lead contact

Further information and requests for resources should be directed to the lead contact, Haiyuan Yu.

## Data and code availability

Data that support the findings of this study are available in the article, supplementary tables, and GitHub repository (https://github.com/haiyuan-yu-lab/3D-Viral-Human and https://github.com/jishnu-lab/ViralHuman3D). Viral-protein structures predicted by AlphaFold are accessible via the web server (https://3d-viralhuman.yulab.org/).

## Acknowledgements

N.R.G. was supported by K99AI175483. L.M.S. was supported by R01AI153396. J.D. was supported by DP2AI164325, U01AI179514, and R01AI170108. H.Y. was supported by SFARI893926, R01EB027895, RM1GM139738, RF1AG082211.

## Author contributions

Conceptualization: L.L., P.G.R., J.D., H.Y.

Methodology: L.L., P.G.R., L.M.S., J.D., H.Y.

Investigation: L.L., P.G.R., and N.S.G.

Formal Analysis: L.L., P.G.R., Y.L., Z.Z., D.X., R.S., N.S.G.

Writing: L.L., P.G.R., J.D., H.Y. with inputs from all authors;

Funding Acquisition: J.D., H.Y.

Resources: R.S., J.D., and H.Y.

Supervision: L.M.S., J.D., and H.Y.

L.L. and P.G.R. contributed equally to the manuscript as co-first authors.

## References

1. Abramson, Josh, Jonas Adler, Jack Dunger, Richard Evans, Tim Green, Alexander Pritzel, Olaf Ronneberger, et al. 2024. “Accurate Structure Prediction of Biomolecular Interactions with AlphaFold 3.” Nature 630 (8016): 493–500.

2. Alqahtani, Aminah, and Meznah Almutairy. 2023. “Evaluating the Performance of Multiple Sequence Alignment Programs with Application to Genotyping SARS-CoV-2 in the Saudi Population.” *Computation (Basel*, Switzerland) 11 (11): 212.

3. Alshareeda, A. T., O. H. Negm, A. R. Green, C. C. Nolan, P. Tighe, N. Albarakati, R. Sultana, S. Madhusudan, I. O. Ellis, and E. A. Rakha. 2015. “KPNA2 Is a Nuclear Export Protein That Contributes to Aberrant Localisation of Key Proteins and Poor Prognosis of Breast Cancer.” British Journal of Cancer 112 (12): 1929–37.

4. Barreiro, Luis B., Meriem Ben-Ali, Hélène Quach, Guillaume Laval, Etienne Patin, Joseph K. Pickrell, Christiane Bouchier, et al. 2009. “Evolutionary Dynamics of Human Toll-like Receptors and Their Different Contributions to Host Defense.” PLoS Genetics 5 (7): e1000562.

5. Basu, Sankar, and Björn Wallner. 2016. “DockQ: A Quality Measure for Protein-Protein Docking Models.” PloS One 11 (8): e0161879.

6. Boutell, Chris, and Roger D. Everett. 2013. “Regulation of Alphaherpesvirus Infections by the ICP0 Family of Proteins.” The Journal of General Virology 94 (Pt 3): 465–81.

7. Bryant, Patrick, and Frank Noé. 2023. “Improved Protein Complex Prediction with AlphaFold-Multimer by Denoising the MSA Profile.” bioRxiv. 10.1101/2023.07.04.547638.

8. Burd, Eileen M. 2003. “Human Papillomavirus and Cervical Cancer.” Clinical Microbiology Reviews 16 (1): 1.

9. Cakir, Merve, Kirsten Obernier, Antoine Forget, and Nevan J. Krogan. 2021. “Target Discovery for Host-Directed Antiviral Therapies: Application of Proteomics Approaches.” *mSystems*, September. 10.1128/msystems.00388-21.

10. Chassey, Benoît de, Laurène Meyniel-Schicklin, Jacky Vonderscher, Patrice André, and Vincent Lotteau. 2014. “Virus-Host Interactomics: New Insights and Opportunities for Antiviral Drug Discovery.” Genome Medicine 6 (11): 1–14.

11. Chhibbar, Prabal, Priyamvada Guha Roy, Munesh K. Harioudh, Daniel J. McGrail, Donghui Yang, Harinder Singh, Reinhard Hinterleitner, et al. 2024. “Uncovering Cell-Type-Specific Immunomodulatory Variants and Molecular Phenotypes in COVID-19 Using Structurally Resolved Protein Networks.” Cell Reports 43 (11): 114930.

12. Cho, Sunglim, Bo Young Kim, Kwangseog Ahn, and Youngsoo Jun. 2013. “The C-Terminal Amino Acid of the MHC-I Heavy Chain Is Critical for Binding to Derlin-1 in Human Cytomegalovirus US11-Induced MHC-I Degradation.” PloS One 8 (8): e72356.

13. Cho, Sunglim, Miriam Lee, and Youngsoo Jun. 2013. “Forced Interaction of Cell Surface Proteins with Derlin-1 in the Endoplasmic Reticulum Is Sufficient to Induce Their Dislocation into the Cytosol for Degradation.” Biochemical and Biophysical Research Communications 430 (2): 787–92.

14. Christiansen, Anders, and Lars Dyrskjøt. 2013. “The Functional Role of the Novel Biomarker Karyopherin α 2 (KPNA2) in Cancer.” Cancer Letters 331 (1): 18–23.

15. Cohen, Paul A., Anjua Jhingran, Ana Oaknin, and Lynette Denny. 2019. “Cervical Cancer.” Lancet 393 (10167): 169–82.

16. Danecek, Petr, Adam Auton, Goncalo Abecasis, Cornelis A. Albers, Eric Banks, Mark A. DePristo, Robert E. Handsaker, et al. 2011. “The Variant Call Format and VCFtools.” *Bioinformatics (Oxford*, England) 27 (15): 2156–58.

17. Desbien, Anthony L., John W. Kappler, and Philippa Marrack. 2009. “The Epstein-Barr Virus Bcl-2 Homolog, BHRF1, Blocks Apoptosis by Binding to a Limited Amount of Bim.” Proceedings of the National Academy of Sciences of the United States of America 106 (14): 5663–68.

18. Dominguez, Cyril, Rolf Boelens, and Alexandre M. J. J. Bonvin. 2003. “HADDOCK: A Protein-Protein Docking Approach Based on Biochemical or Biophysical Information.” Journal of the American Chemical Society 125 (7): 1731–37.

19. Doorbar, John, Wim Quint, Lawrence Banks, Ignacio G. Bravo, Mark Stoler, Tom R. Broker, and Margaret A. Stanley. 2012. “The Biology and Life-Cycle of Human Papillomaviruses.” Vaccine 30 Suppl 5 (November):F55–70.

20. Drews, Camille M., Samuel Case, and Scott B. Vande Pol. 2019. “E6 Proteins from High-Risk HPV, Low-Risk HPV, and Animal Papillomaviruses Activate the Wnt/β-Catenin Pathway through E6AP-Dependent Degradation of NHERF1.” PLoS Pathogens 15 (4): e1007575.

21. Evans, Richard, Michael O’Neill, Alexander Pritzel, Natasha Antropova, Andrew Senior, Tim Green, Augustin Žídek, et al. 2022. “Protein Complex Prediction with AlphaFold-Multimer.” bioRxiv. 10.1101/2021.10.04.463034.

22. Fang, Run, Qifei Jiang, Xiang Zhou, Chenguang Wang, Yukun Guan, Jianli Tao, Jianzhong Xi, Ji-Ming Feng, and Zhengfan Jiang. 2017. “MAVS Activates TBK1 and IKKε through TRAFs in NEMO Dependent and Independent Manner.” PLoS Pathogens 13 (11): e1006720.

23. Fätkenheuer, Gerd, Anton L. Pozniak, Margaret A. Johnson, Andreas Plettenberg, Schlomo Staszewski, Andy I. M. Hoepelman, Michael S. Saag, et al. 2005. “Efficacy of Short-Term Monotherapy with Maraviroc, a New CCR5 Antagonist, in Patients Infected with HIV-1.” Nature Medicine 11 (11): 1170–72.

24. Ferrari, Mathieu, Leila Mekkaoui, F. Tudor Ilca, Zulaikha Akbar, Reyisa Bughda, Katarina Lamb, Katarzyna Ward, et al. 2021. “Characterization of a Novel ACE2-Based Therapeutic with Enhanced rather than Reduced Activity against SARS-CoV-2 Variants.” Journal of Virology 95 (19): e0068521.

25. Franzosa, Eric A., and Yu Xia. 2011. “Structural Principles within the Human-Virus Protein-Protein Interaction Network.” Proceedings of the National Academy of Sciences of the United States of America 108 (26): 10538–43.

26. Fumagalli, Matteo, Manuela Sironi, Uberto Pozzoli, Anna Ferrer-Admetlla, Linda Pattini, and Rasmus Nielsen. 2011. “Signatures of Environmental Genetic Adaptation Pinpoint Pathogens as the Main Selective Pressure through Human Evolution.” PLoS Genetics 7 (11): e1002355.

27. Gokhale, Nandan S., Russell K. Sam, Kim Somfleth, Matthew G. Thompson, Daphnée M. Marciniak, Julian R. Smith, Emmanuelle Genoyer, et al. 2024. “Cellular RNA Interacts with MAVS to Promote Antiviral Signaling.” *Science (New York*, N.Y.) 386 (6728): eadl0429.

28. Gordon, David E., Joseph Hiatt, Mehdi Bouhaddou, Veronica V. Rezelj, Svenja Ulferts, Hannes Braberg, Alexander S. Jureka, et al. 2020. “Comparative Host-Coronavirus Protein Interaction Networks Reveal Pan-Viral Disease Mechanisms.” *Science (New York*, N.Y.) 370 (6521): eabe9403.

29. Grimmel, M., E-M De Villiers, Ch Neumann, M. Pawlita, and H. zur Hausen. 1988. “Characterization of a New Human Papillomavirus (HPV 41) from Disseminated Warts and Detection of Its DNA in Some Skin Carcinomas.” International Journal of Cancer 41 (1): 5– 9.

30. Guven-Maiorov, Emine, Asma Hakouz, Sukejna Valjevac, Ozlem Keskin, Chung-Jung Tsai, Attila Gursoy, and Ruth Nussinov. 2020. “HMI-PRED: A Web Server for Structural Prediction of Host-Microbe Interactions Based on Interface Mimicry.” Journal of Molecular Biology 432 (11): 3395–3403.

31. Holmes, Edward C. 2003. “Molecular Clocks and the Puzzle of RNA Virus Origins.” Journal of Virology 77 (7): 3893–97.

32. Jumper, John, Richard Evans, Alexander Pritzel, Tim Green, Michael Figurnov, Olaf Ronneberger, Kathryn Tunyasuvunakool, et al. 2021. “Highly Accurate Protein Structure Prediction with AlphaFold.” Nature 596 (7873): 583–89.

33. Karlsson, Elinor K., Dominic P. Kwiatkowski, and Pardis C. Sabeti. 2014. “Natural Selection and Infectious Disease in Human Populations.” Nature Reviews. Genetics 15 (6): 379–93.

34. Katoh, Kazutaka, Kazuharu Misawa, Kei-Ichi Kuma, and Takashi Miyata. 2002. “MAFFT: A Novel Method for Rapid Multiple Sequence Alignment Based on Fast Fourier Transform.” Nucleic Acids Research 30 (14): 3059–66.

35. Ketata, Mohamed Amine, Cedrik Laue, Ruslan Mammadov, Hannes Stärk, Menghua Wu, Gabriele Corso, Céline Marquet, Regina Barzilay, and Tommi S. Jaakkola. 2023. “DiffDock-PP: Rigid Protein-Protein Docking with Diffusion Models.” arXiv [q-bio.BM]. 10.48550/ARXIV.2304.03889.

36. Lan, Jun, Jiwan Ge, Jinfang Yu, Sisi Shan, Huan Zhou, Shilong Fan, Qi Zhang, et al. 2020. “Structure of the SARS-CoV-2 Spike Receptor-Binding Domain Bound to the ACE2 Receptor.” Nature 581 (7807): 215–20.

37. Lasso, Gorka, Barry Honig, and Sagi D. Shapira. 2021. “A Sweep of Earth’s Virome Reveals Host-Guided Viral Protein Structural Mimicry and Points to Determinants of Human Disease.” Cell Systems 12 (1): 82–91.e3.

38. Lasso, Gorka, Sandra V. Mayer, Evandro R. Winkelmann, Tim Chu, Oliver Elliot, Juan Angel Patino-Galindo, Kernyu Park, Raul Rabadan, Barry Honig, and Sagi D. Shapira. 2019. “A Structure-Informed Atlas of Human-Virus Interactions.” Cell 178 (6): 1526–41.e16.

39. Le Roux, Lucia G., and Junona Moroianu. 2003. “Nuclear Entry of High-Risk Human Papillomavirus Type 16 E6 Oncoprotein Occurs via Several Pathways.” Journal of Virology 77 (4): 2330–37.

40. Lewontin, R. C., and J. Krakauer. 1973. “Distribution of Gene Frequency as a Test of the Theory of the Selective Neutrality of Polymorphisms.” Genetics 74 (1): 175–95.

41. Lilley, Brendan N., and Hidde L. Ploegh. 2004. “A Membrane Protein Required for Dislocation of Misfolded Proteins from the ER.” Nature 429 (6994): 834–40.

42. Lin, Zeming, Halil Akin, Roshan Rao, Brian Hie, Zhongkai Zhu, Wenting Lu, Nikita Smetanin, et al. 2023. “Evolutionary-Scale Prediction of Atomic-Level Protein Structure with a Language Model.” *Science (New York*, N.Y.) 379 (6637): 1123–30.

43. Liu, Siqi, Jueqi Chen, Xin Cai, Jiaxi Wu, Xiang Chen, You-Tong Wu, Lijun Sun, and Zhijian J. Chen. 2013. “MAVS Recruits Multiple Ubiquitin E3 Ligases to Activate Antiviral Signaling Cascades.” eLife 2 (August):e00785.

44. Liu, Xiaoyang, Senthilkumar Palaniyandi, Iowis Zhu, Jin Tang, Weizhong Li, Xiaoling Wu, Susan Park Ochsner, C. David Pauza, Jeffrey I. Cohen, and Xiaoping Zhu. 2019. “Human Cytomegalovirus Evades Antibody-Mediated Immunity through Endoplasmic Reticulum-Associated Degradation of the FcRn Receptor.” Nature Communications 10 (1): 1–19.

45. Liu, Xueqiao, Katherine Fitzgerald, Evelyn Kurt-Jones, Robert Finberg, and David M. Knipe. 2008. “Herpesvirus Tegument Protein Activates NF-kappaB Signaling through the TRAF6 Adaptor Protein.” Proceedings of the National Academy of Sciences of the United States of America 105 (32): 11335–39.

46. Longworth, Michelle S., and Laimonis A. Laimins. 2004. “Pathogenesis of Human Papillomaviruses in Differentiating Epithelia.” Microbiology and Molecular Biology Reviews: MMBR 68 (2): 362–72.

47. Lupo, Umberto, Damiano Sgarbossa, and Anne-Florence Bitbol. 2024. “Pairing Interacting Protein Sequences Using Masked Language Modeling.” Proceedings of the National Academy of Sciences of the United States of America 121 (27): e2311887121.

48. Mantovani, F., and L. Banks. 2001. “The Human Papillomavirus E6 Protein and Its Contribution to Malignant Progression.” Oncogene 20 (54): 7874–87.

49. Martel, Catherine de, Martyn Plummer, Jerome Vignat, and Silvia Franceschi. 2017. “Worldwide Burden of Cancer Attributable to HPV by Site, Country and HPV Type.” International Journal of Cancer. Journal International Du Cancer 141 (4): 664–70.

50. Ma, Shouzhi, and Xiaohang Zhao. 2014. “KPNA2 Is a Promising Biomarker Candidate for Esophageal Squamous Cell Carcinoma and Correlates with Cell Proliferation.” Oncology Reports 32 (4): 1631–37.

51. Mesplède, Thibault, David Gagnon, Fanny Bergeron-Labrecque, Ibrahim Azar, Hélène Sénéchal, François Coutlée, and Jacques Archambault. 2012. “p53 Degradation Activity, Expression, and Subcellular Localization of E6 Proteins from 29 Human Papillomavirus Genotypes.” Journal of Virology 86 (1): 94–107.

52. Meyer, Michael J., Juan Felipe Beltrán, Siqi Liang, Robert Fragoza, Aaron Rumack, Jin Liang, Xiaomu Wei, and Haiyuan Yu. 2018. “Interactome INSIDER: A Structural Interactome Browser for Genomic Studies.” Nature Methods 15 (2): 107–14.

53. Meyer, Michael J., Philip Geske, and Haiyuan Yu. 2016. “BISQUE: Locus– and Variant-Specific Conversion of Genomic, Transcriptomic and Proteomic Database Identifiers.” *Bioinformatics (Oxford*, England) 32 (10): 1598–1600.

54. Meyerson, Nicholas R., Paul A. Rowley, Christina H. Swan, Dona T. Le, Gregory K. Wilkerson, and Sara L. Sawyer. 2014. “Positive Selection of Primate Genes That Promote HIV-1 Replication.” Virology 454–455 (April):291–98.

55. Mihalič, Filip, Leandro Simonetti, Girolamo Giudice, Marie Rubin Sander, Richard Lindqvist, Marie Berit Akpiroro Peters, Caroline Benz, et al. 2023. “Large-Scale Phage-Based Screening Reveals Extensive Pan-Viral Mimicry of Host Short Linear Motifs.” Nature Communications 14 (1): 2409.

56. Mukherjee, Srayanta, and Yang Zhang. 2009. “MM-Align: A Quick Algorithm for Aligning Multiple-Chain Protein Complex Structures Using Iterative Dynamic Programming.” Nucleic Acids Research 37 (11): e83.

57. “NCBI Virus.” n.d. Accessed February 4, 2025. https://www.ncbi.nlm.nih.gov/labs/virus/vssi/#/.

58. Oda, Kosuke, Yasuyuki Matoba, Masanori Sugiyama, and Takemasa Sakaguchi. 2021. “Structural Insight into the Interaction of Sendai Virus C Protein with Alix to Stimulate Viral Budding.” Journal of Virology 95 (19): e0081521.

59. Oh, Stephen T., Michelle S. Longworth, and Laimonis A. Laimins. 2004. “Roles of the E6 and E7 Proteins in the Life Cycle of Low-Risk Human Papillomavirus Type 11.” Journal of Virology 78 (5): 2620–26.

60. Pal, Asmita, and Rita Kundu. 2019. “Human Papillomavirus E6 and E7: The Cervical Cancer Hallmarks and Targets for Therapy.” Frontiers in Microbiology 10:3116.

61. Perez, Gerardo, Galt P. Barber, Anna Benet-Pages, Jonathan Casper, Hiram Clawson, Mark Diekhans, Clay Fischer, et al. 2025. “The UCSC Genome Browser Database: 2025 Update.” Nucleic Acids Research 53 (D1): D1243–49.

62. Sahni, Nidhi, Song Yi, Mikko Taipale, Juan I. Fuxman Bass, Jasmin Coulombe-Huntington, Fan Yang, Jian Peng, et al. 2015. “Widespread Macromolecular Interaction Perturbations in Human Genetic Disorders.” Cell 161 (3): 647–60.

63. Saraiya, Mona, Elizabeth R. Unger, Trevor D. Thompson, Charles F. Lynch, Brenda Y. Hernandez, Christopher W. Lyu, Martin Steinau, et al. 2015. “US Assessment of HPV Types in Cancers: Implications for Current and 9-Valent HPV Vaccines.” Journal of the National Cancer Institute 107 (6): djv086.

64. Schiffman, Mark, John Doorbar, Nicolas Wentzensen, Silvia de Sanjosé, Carole Fakhry, Bradley J. Monk, Margaret A. Stanley, and Silvia Franceschi. 2016. “Carcinogenic Human Papillomavirus Infection.” Nature Reviews. Disease Primers 2 (December):16086.

65. Shoemaker, Robert H., Reynold A. Panettieri Jr, Steven K. Libutti, Howard S. Hochster, Norman R. Watts, Paul T. Wingfield, Philipp Starkl, et al. 2022. “Development of an Aerosol Intervention for COVID-19 Disease: Tolerability of Soluble ACE2 (APN01) Administered via Nebulizer.” PloS One 17 (7): e0271066.

66. Sironi, Manuela, Rachele Cagliani, Diego Forni, and Mario Clerici. 2015. “Evolutionary Insights into Host-Pathogen Interactions from Mammalian Sequence Data.” Nature Reviews. Genetics 16 (4): 224–36.

67. Smith, Miles C., Chris Boutell, and David J. Davido. 2011. “HSV-1 ICP0: Paving the Way for Viral Replication.” Future Virology 6 (4): 421–29.

68. Syrjänen, S. M., and K. J. Syrjänen. 1999. “New Concepts on the Role of Human Papillomavirus in Cell Cycle Regulation.” Annals of Medicine 31 (3): 175–87.

69. Thomas, M., and L. Banks. 1999. “Human Papillomavirus (HPV) E6 Interactions with Bak Are Conserved amongst E6 Proteins from High and Low Risk HPV Types.” The Journal of General Virology 80 (Pt 6) (6): 1513–17.

70. Thomas, Miranda, Richard Laura, Karin Hepner, Ernesto Guccione, Charles Sawyers, Laurence Lasky, and Lawrence Banks. 2002. “Oncogenic Human Papillomavirus E6 Proteins Target the MAGI-2 and MAGI-3 Proteins for Degradation.” Oncogene 21 (33): 5088–96.

71. Tsuchiya, Yuko, Yu Yamamori, and Kentaro Tomii. 2022. “Protein-Protein Interaction Prediction Methods: From Docking-Based to AI-Based Approaches.” Biophysical Reviews 14 (6): 1341–48.

72. Underbrink, Michael P., Crystal Dupuis, Jia Wang, and Stephen K. Tyring. 2016. “E6 Proteins from Low-Risk Human Papillomavirus Types 6 and 11 Are Able to Protect Keratinocytes from Apoptosis via Bak Degradation.” The Journal of General Virology 97 (3): 715–24.

73. Van Blerkom, Linda M. 2003. “Role of Viruses in Human Evolution.” American Journal of Physical Anthropology Suppl 37 (S37): 14–46.

74. Vanni, Emilia, Derek Gatherer, Lily Tong, Roger D. Everett, and Chris Boutell. 2012. “Functional Characterization of Residues Required for the Herpes Simplex Virus 1 E3 Ubiquitin Ligase ICP0 to Interact with the Cellular E2 Ubiquitin-Conjugating Enzyme UBE2D1 (UbcH5a).” Journal of Virology 86 (11): 6323–33.

75. Wang, Xiujuan, Xiaomu Wei, Bram Thijssen, Jishnu Das, Steven M. Lipkin, and Haiyuan Yu. 2012. “Three-Dimensional Reconstruction of Protein Networks Provides Insight into Human Genetic Disease.” Nature Biotechnology 30 (2): 159–64.

76. Watt, Pauline J. van der, Ellen Ngarande, and Virna D. Leaner. 2011. “Overexpression of Kpnβ1 and Kpnα2 Importin Proteins in Cancer Derives from Deregulated E2F Activity.” PloS One 6 (11): e27723.

77. Wierbowski, Shayne D., Siqi Liang, Yuan Liu, You Chen, Shagun Gupta, Nicole M. Andre, Steven M. Lipkin, Gary R. Whittaker, and Haiyuan Yu. 2021. “A 3D Structural SARS-CoV-2–human Interactome to Explore Genetic and Drug Perturbations.” Nature Methods 18 (11): 1477–88.

78. Wright, Sewall. 1965. “The Interpretation of Population Structure by F-Statistics with Special Regard to Systems of Mating.” Evolution; International Journal of Organic Evolution 19 (3): 395.

79. Xiong, Dapeng, Yunguang Qiu, Junfei Zhao, Yadi Zhou, Dongjin Lee, Shobhita Gupta, Mateo Torres, et al. 2024. “A Structurally Informed Human Protein-Protein Interactome Reveals Proteome-Wide Perturbations Caused by Disease Mutations.” *Nature Biotechnology*, October, 1–15.

80. Xu, Liang-Guo, Yan-Yi Wang, Ke-Jun Han, Lian-Yun Li, Zhonghe Zhai, and Hong-Bing Shu. 2005. “VISA Is an Adapter Protein Required for Virus-Triggered IFN-Beta Signaling.” Molecular Cell 19 (6): 727–40.

81. Yim, Eun-Kyoung, and Jong-Sup Park. 2005. “The Role of HPV E6 and E7 Oncoproteins in HPV-Associated Cervical Carcinogenesis.” Cancer Research and Treatment: Official Journal of Korean Cancer Association 37 (6): 319–24.

82. Yi, Sang Ah, Dong Hoon Lee, Go Woon Kim, Hyun-Wook Ryu, Jong Woo Park, Jaecheol Lee, Jihoon Han, et al. 2020. “HPV-Mediated Nuclear Export of HP1γ Drives Cervical Tumorigenesis by Downregulation of p53.” Cell Death and Differentiation 27 (9): 2537–51.

83. Zhang, Junjie, Jun Zhao, Simin Xu, Junhua Li, Shanping He, Yi Zeng, Linshen Xie, et al. 2018. “Species-Specific Deamidation of cGAS by Herpes Simplex Virus UL37 Protein Facilitates Viral Replication.” Cell Host & Microbe 24 (2): 234–48.e5.

84. Zhao, Jun, Yi Zeng, Simin Xu, Jie Chen, Guobo Shen, Caiqun Yu, David Knipe, et al. 2016. “A Viral Deamidase Targets the Helicase Domain of RIG-I to Block RNA-Induced Activation.” Cell Host & Microbe 20 (6): 770–84.

85. Zhu, Wensi, Aditi Shenoy, Petras Kundrotas, and Arne Elofsson. 2023. “Evaluation of AlphaFold-Multimer Prediction on Multi-Chain Protein Complexes.” Bioinformatics 39 (7). 10.1093/bioinformatics/btad424.

