## Supplementary figures and images for "Comprehensive Atomic-Scale 3D Viral-Host Protein Interactomes Enable Dissection of Key Mechanisms and Evolutionary Processes Underlying Viral Pathogenesis"

### Supplementary Figures

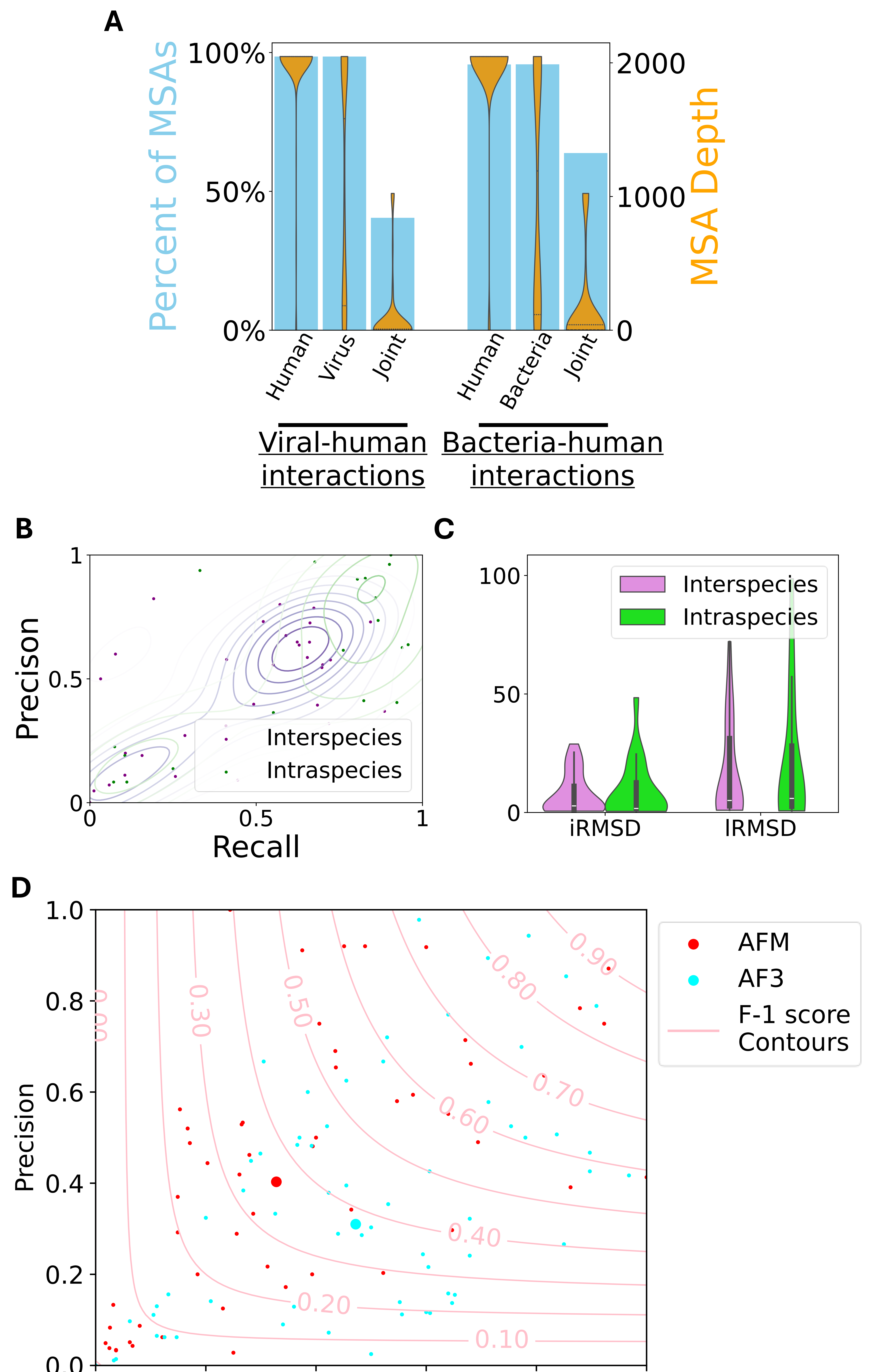

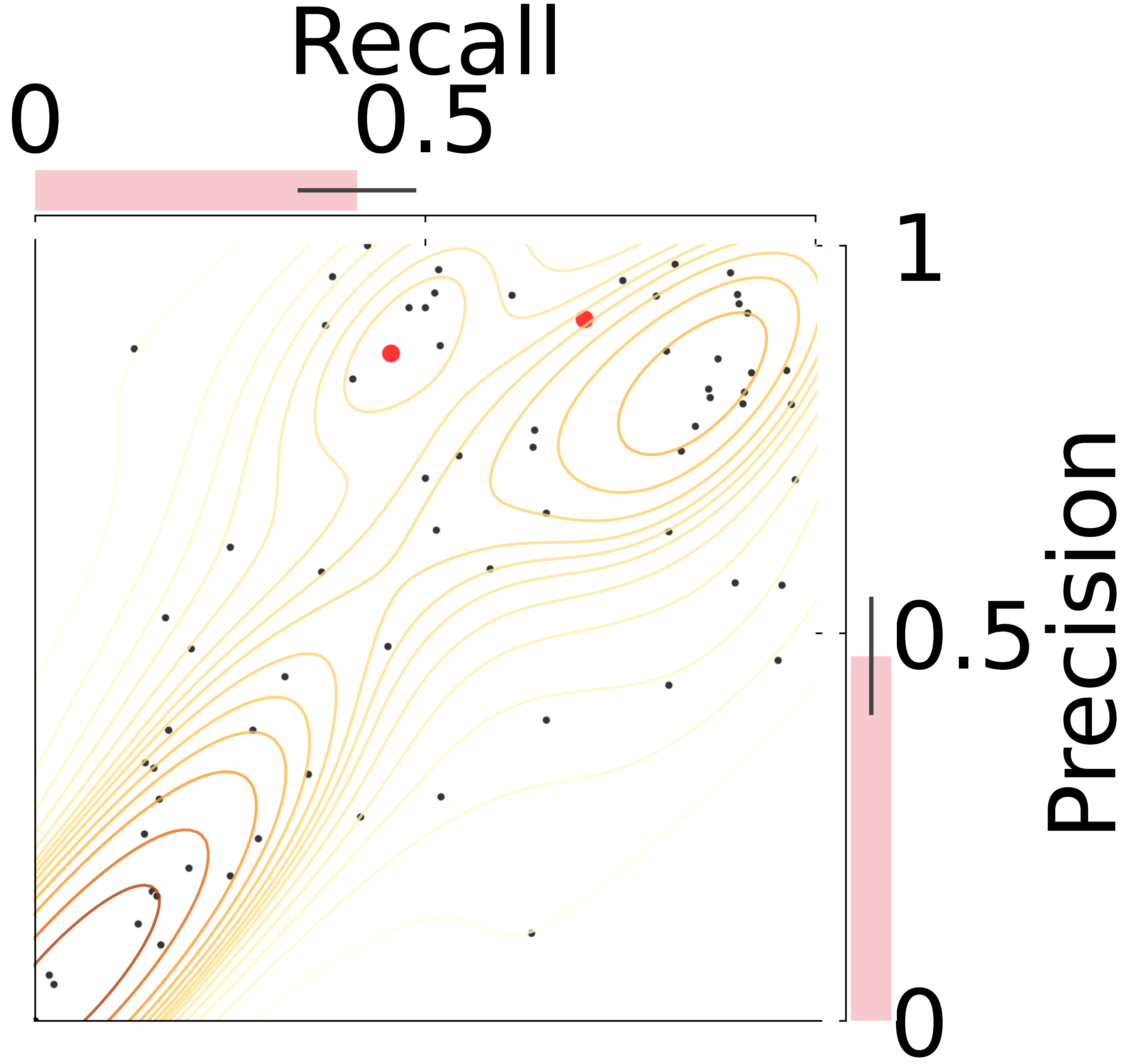

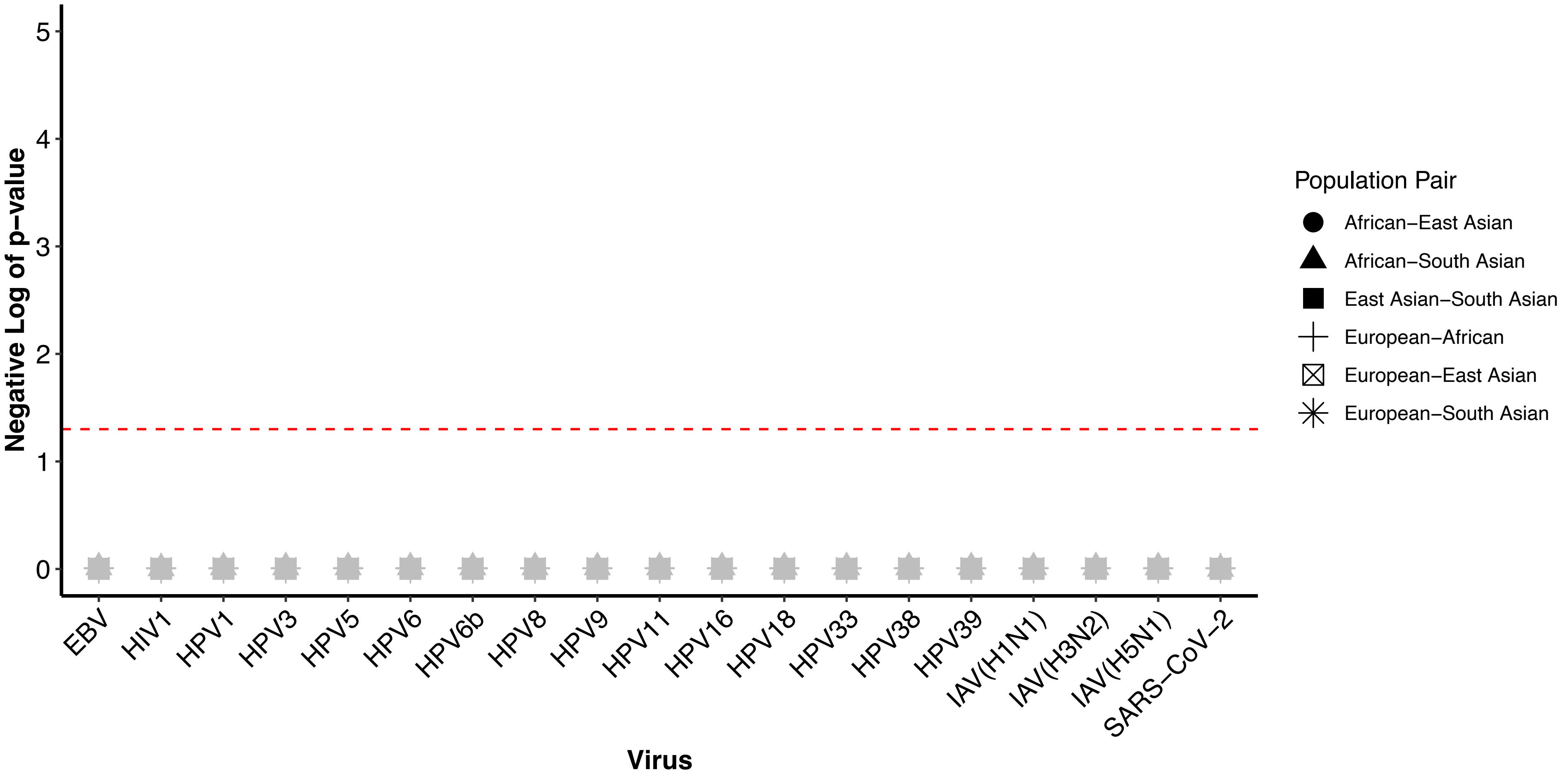

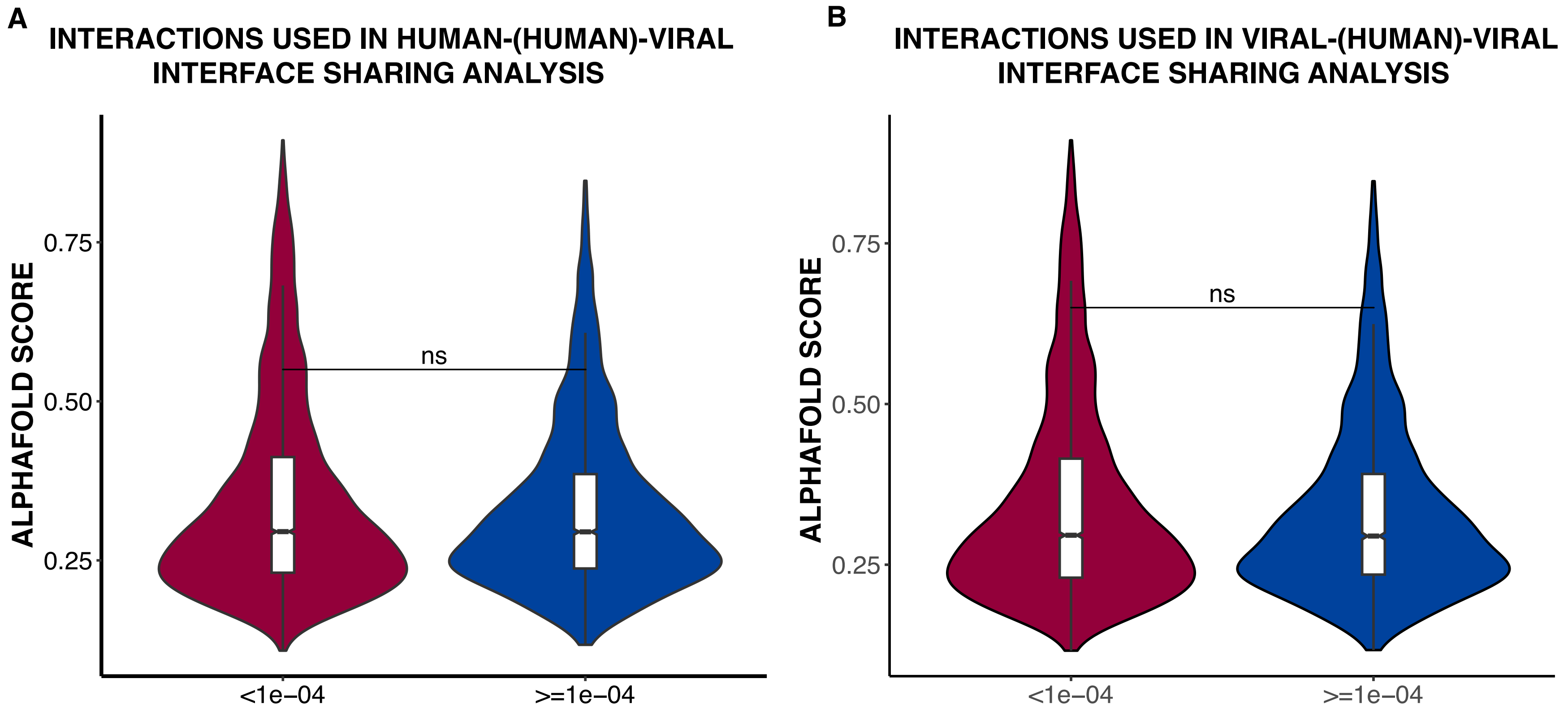

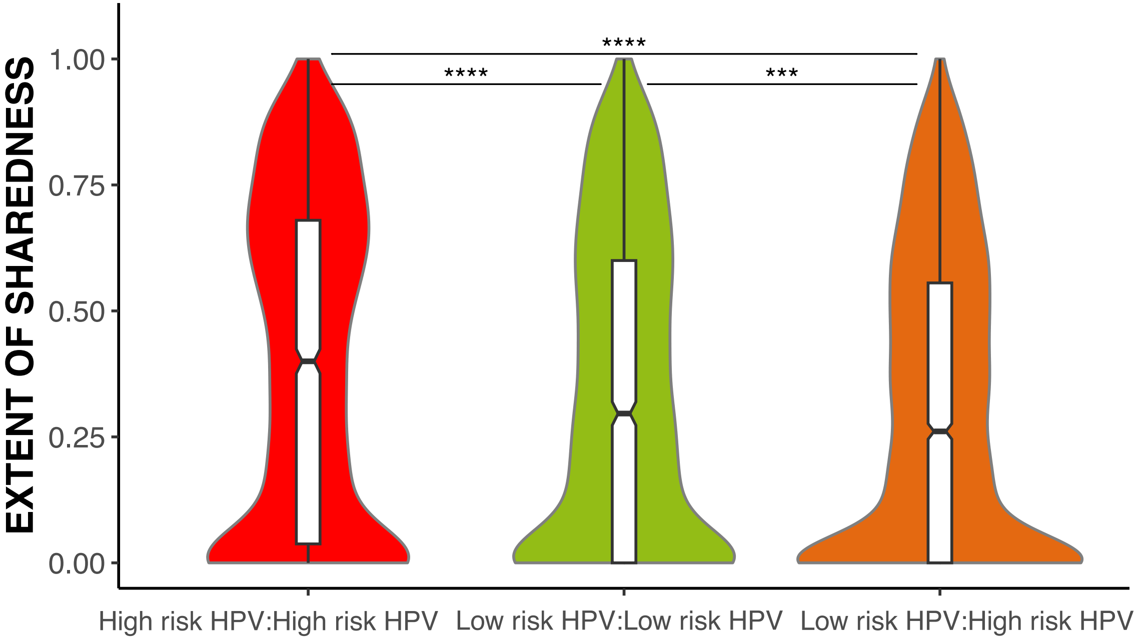

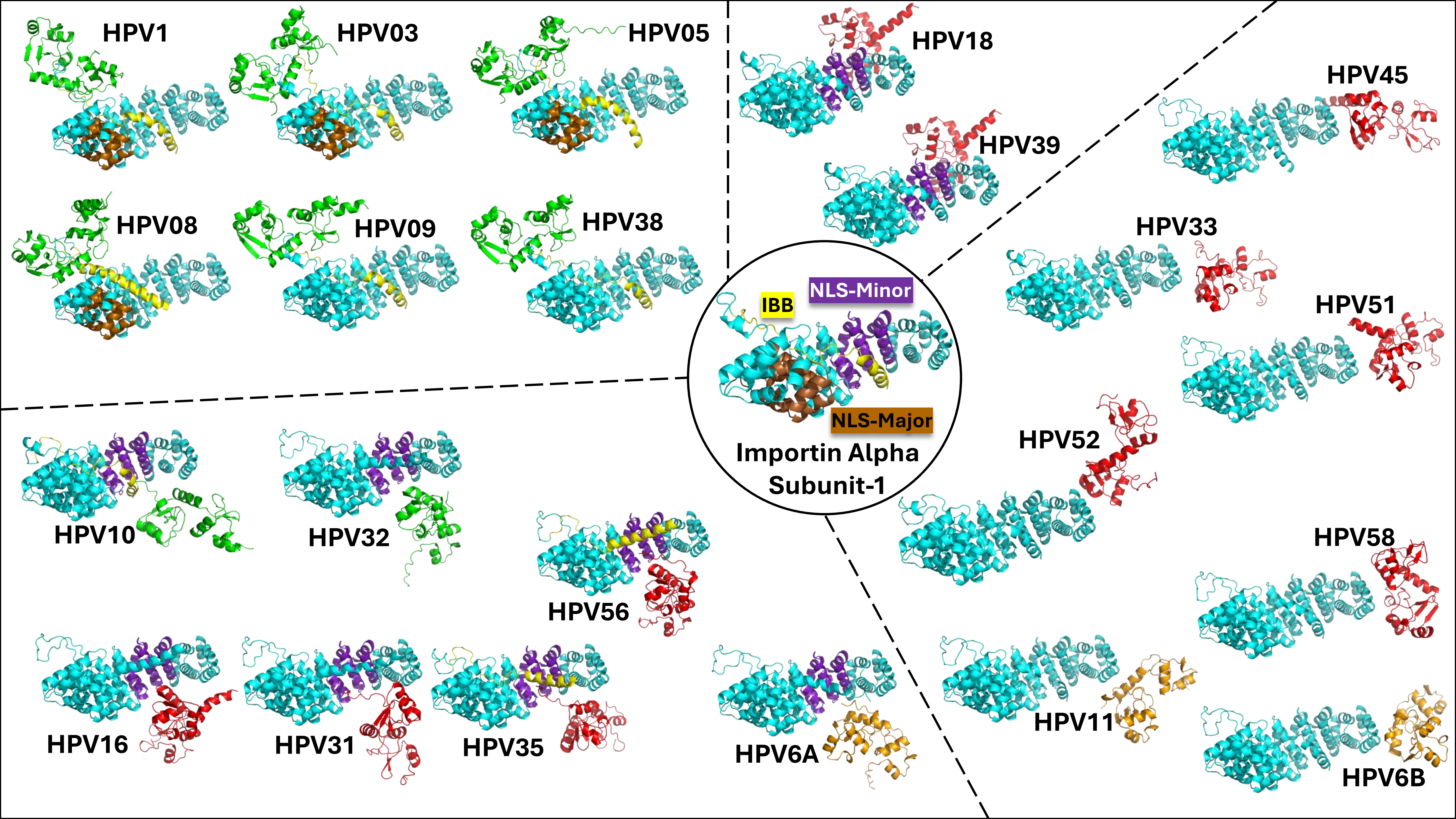

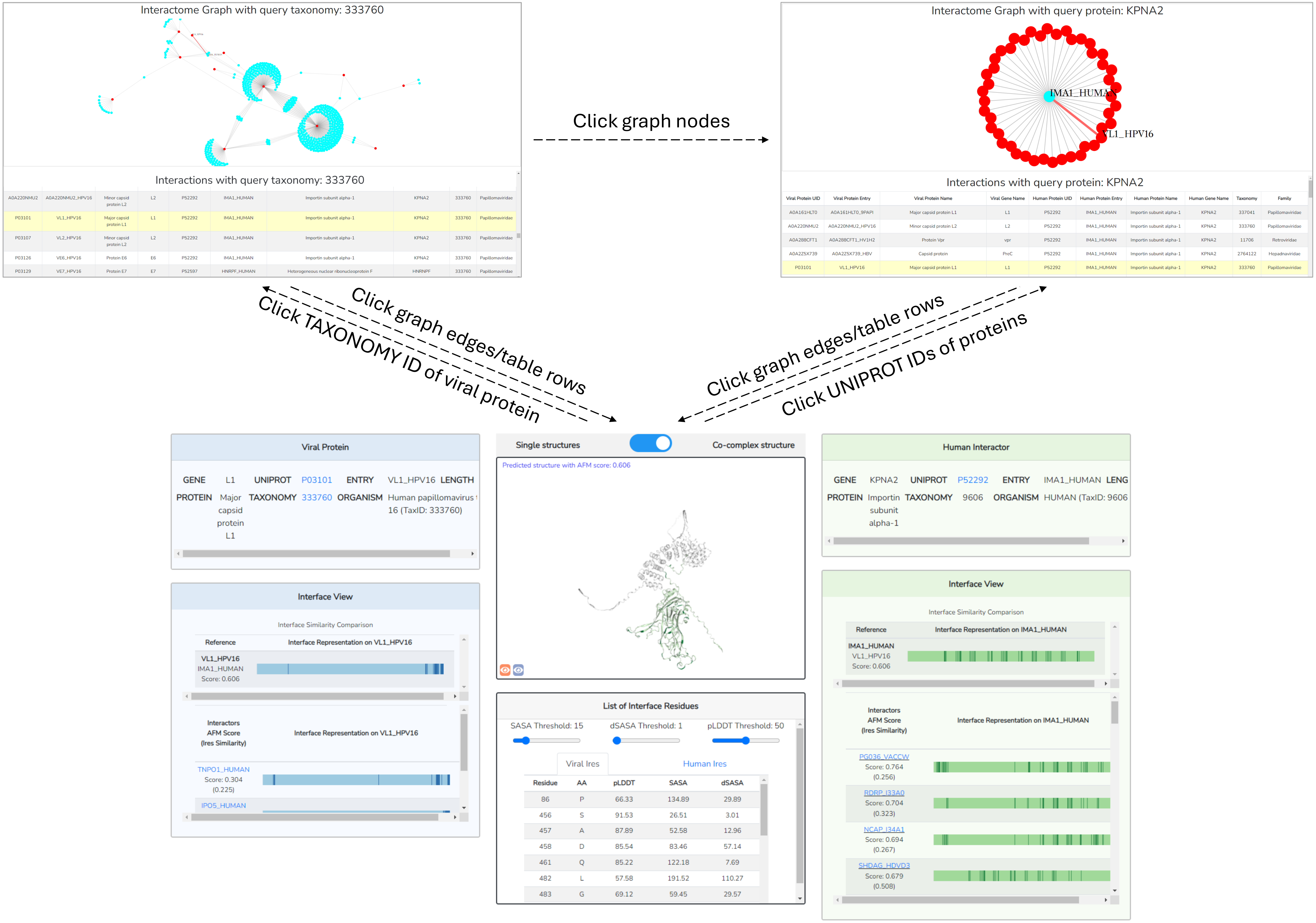
